# Rock inhibitors target *SRSF2* leukemia by disrupting cell mitosis and nuclear morphology

**DOI:** 10.1101/2022.02.15.479934

**Authors:** M Su, T Fleisher, I Grosheva, M Bokstad Horev, M Olszewska, H Barr, A Plotnikov, S Carvalho, Y Moskovich, MD Minden, N Chapal-Ilani, EP Papapetrou, N Dezorella, T Cheng, N Kaushansky, B Geiger, LI Shlush

## Abstract

Spliceosome machinery mutations are common early mutations in myeloid malignancies, however effective targeted therapies against them are still lacking. In the current study, we used an *in vitro* high-throughput drug screen among four different isogenic cell lines and identified ROCK inhibitors (ROCKi) as selective inhibitors of *SRSF2* mutants. ROCKi targeted *SRSF2* Mut primary human samples in a xenografts model and were not toxic to mice nor human cells. ROCKi induced mitotic catastrophe through their apparent effects on microtubules and nuclear organization. Transmission electron microscopy revealed that *SRSF2* mutations induce deep nuclear indentation and segmentation, driven by microtubule-rich cytoplasmic intrusions, which were exacerbated by ROCKi. The severe nuclear deformation driven by the combination of *SRSF2* Mut and ROCKi prevent cells from completing mitosis. These findings shed light on new ways to target *SRSF2* and on the role of the microtubule system in *SRSF2* Mut cells.

## INTRODUCTION

Splicing machinery mutations (SMMs) can be frequently identified in various myeloid malignancies (acute myeloid leukemia (AML), myeloid dysplastic syndrome (MDS), chronic myelomonocytic leukaemia (CMML), myeloproliferative neoplasms (MPN)). Furthermore, it has been discovered they can be identified among healthy individuals as part of age-related clonal hematopoiesis (CH)^1^. The presence of *SRSF2* or *U2AF1* mutations among healthy individuals defines high-risk individuals of progressing to overt AML^1,2^. The fact that *SRSF2*/*U2AF1* mutations are the first events in a high-risk leukemogenic process highlights their potential as targets for preventing leukemia. While screening programs for early diagnosis of AML are still not in place, we propose that the next crucial step towards preventive therapy is the creation of an arsenal of safe interventions for targeting SMMs, that could be administrated to high-risk individuals years before leukemia is diagnosed. To achieve such goal, we have focused our efforts on identifying safe therapies that target hematopoietic cells carrying *SRSF2* mutations.

SMMs are commonly heterozygous, require a functioning wild type (WT) allele, and are less tolerant to pharmacological inhibition of splicing^3^. This rationale was successfully tested for one such splicing inhibitor, E7107, which decreased leukemia burden in mice ^4^. However, phase-I clinical studies in which E7107 was administered to patients with myeloid malignancies were prematurely discontinued due to severe adverse effects^5^. H3B-8800, an orally available analog of E7107, has been safely tested on MDS, AML, and CMML patients, but did not induce partial/complete response^6^. Others attempted to target SMMs by identifying specific pathogenic aberrant isoforms^7,8^. Another approach was a functional CRISPR screen to identify pathways crucial to cells carrying SRSF2 mutations^9^.

So far, any of these approaches demonstrated clinical benefit. Altogether, emphasizing that the need for anti-*SRSF2* Mut therapies still exists. Another approach for targeting such a mutation is understanding its pathological consequences and eliciting a synthetic lethal response. In this regard, it is essential to realize that at the cellular level, *SRSF2* mutations are causing more than just mis-splicing errors. Previous data demonstrated that mutated *SRSF2* induced R-loops, DNA replication stress, and G2M cell cycle arrest ^10^.

In this study, we aimed at performing a high-throughput drug screening (HTDS) on *SRSF2* mutated cells and faced the task of establishing an appropriate model. First, cell lines with inherent *SRSF2* Mut are lacking. Second, human *SRSF2* mutated samples are highly heterogeneous and may present with diverse genetic backgrounds. It remains unclear whether the appropriate control cells should be, e.g., *SRSF2* WT AML lines or WT hematopoietic stem and progenitor cells (HSPCs). On top of that, *SRSF2* Mut is common mainly among elderly males^11^, adding to the complexity of choosing the most suitable model for such a screening. Altogether, to perform HTDS, we created five *SRSF2* isogenic cell lines. In this way, we had sufficient background complexity but with well-matched isogenic WT controls.

Here we report the discovery of Rho-associated coiled-coil containing protein kinase inhibitors (ROCKi), which can inhibit *SRSF2* mutated AML cells *in vitro*, *in vivo,* and in primary AML samples. ROCKi enhanced the defective cytoskeleton system, nuclear deformation and segmentation of *SRSF2* mutated cells, and caused mitotic catastrophe, eventually leading to cell death.

## METHODS

### Cell lines

*SRSF2* P95H mutation were introduced with CRISPR/Cas9 using sgRNA and ssODN in K562, OCI-AML2, OCI-AML3, MOLM-14 and MARIMO cell lines. Single-cells were sorted into 96 well plate and cultured for 2-4 weeks and genotyped.

### RNA-seq

RNA seq of all the cell lines are extracted with Qiuck-RNA MagBead (Zymo, R2132). Sequencing Libraries were prepared using INCPM mRNA Seq. NovaSeq 200 cycles reads were sequenced on 2 lane(s) of an Illumina novaseq. The output was ^~^54 million reads per sample. We used rMATS v4.0.2 to assess the differential splicing landscape embedded in RNA-seq data, with cut off FDR<0.05 and ΔPSI >0.1 (10%).

### Cell viability assay

Compounds were dispensed in 384-well plates using an ECHO® 555 liquid handler (Labcyte) and sealed. Cells were incubated for 48 hours. CellTiter-Glo® (G7572, Promega) was dispensed as per the manufacturer’s instructions. The luminescence signal was then measured using a PHERAstar® FSX (BMG Labtech) and results were analyzed using Genedata Screener®. The viability of treated cells was normalized to a vehicle control.

### In vivo

NSG mice (n=5-10/sample) were injected with half million to five million CD3 depleted AML cells from the peripheral blood of six patients with *SRSF2* mutated AML intra-femorally. On day 35 the animals were randomized to RKI-1447 or a carrier control. RKI-1447 is given i.p. 50mg/kg. On day 56 mice were sacrificed and analyzed for human engraftment by flow cytometry.

### Immunofluorescent cell labeling and imaging

Cells were labeled with anti-α-tubulin anti-Lamin B1. Alexa-488 conjugated secondary antibody was used. Cells were imaged on an inverted Olympus confocal FluoView FV1000 IX81 confocal laser-scanning microscope and an inverted Leica SP8 STED3X confocal microscope. The images were processed by Imaris and ImageJ software.

### Transmission electron microscopy

Cells were fixed, postfixed, stained with 2% uranyl acetate in double distilled water for 1 hour, dehydrated and embedded in epoxy resin. Ultrathin sections (70 nm) were obtained with a Leica EMUC7 ultramicrotome and transferred to 200 mesh copper transmission electron microscopy grids (SPI). Grids were stained with lead citrate and examined with a Tecnai SPIRIT transmission electron microscope (Thermo Fisher Scientific). Digital electron micrographs were acquired with a bottom-mounted Gatan OneView camera.

## RESULTS

### *In vitro* High Throughput Drug Screen Reveals ROCK inhibitors as Potent Therapeutic Drug against SRSF2 Mut Cells

In the current study, we adopted an unbiased approach and conducted an HTDS using three commercially available compound libraries. Towards this goal, we introduced *SRSF2* (P95H mutation) by the CRISPR/CAS9 system into multiple hematopoietic cell lines (MOLM14, K562, MARIMO, OCI-AML2, and OCI-AML3) (**Supplementary Table 1**). *SRSF2* mutations lead to a specific aberrant splicing pattern, dominated by increased exon skipping (SE)^12^. To validate that the *SRSF2* Mut we introduced replicates the aberrant splicing patterns observed in primary AMLs, we performed RNA sequencing and splicing analysis on our isogenic cell lines and samples from the Beat-AML study^13^ with rMATS 4.0.2^14^ (**Fig. 1a, Supplementary Table 2**). We found many aberrant SE events in all of our *SRSF2* Mut lines (**Fig. 1a**). Furthermore, the overlap between splice variants found in the Beat-AML data and the five isogenic SRSF2 Mut cell lines was statistically significant (p< 10^−6^, **Fig.1b, Supplementary Table 3**). These results suggested that the isogenic *SRSF2* Mut cell lines successfully recapitulated primary *SRSF2* Mut AML aberrant splicing and were a valid model.

**Figure 1:**
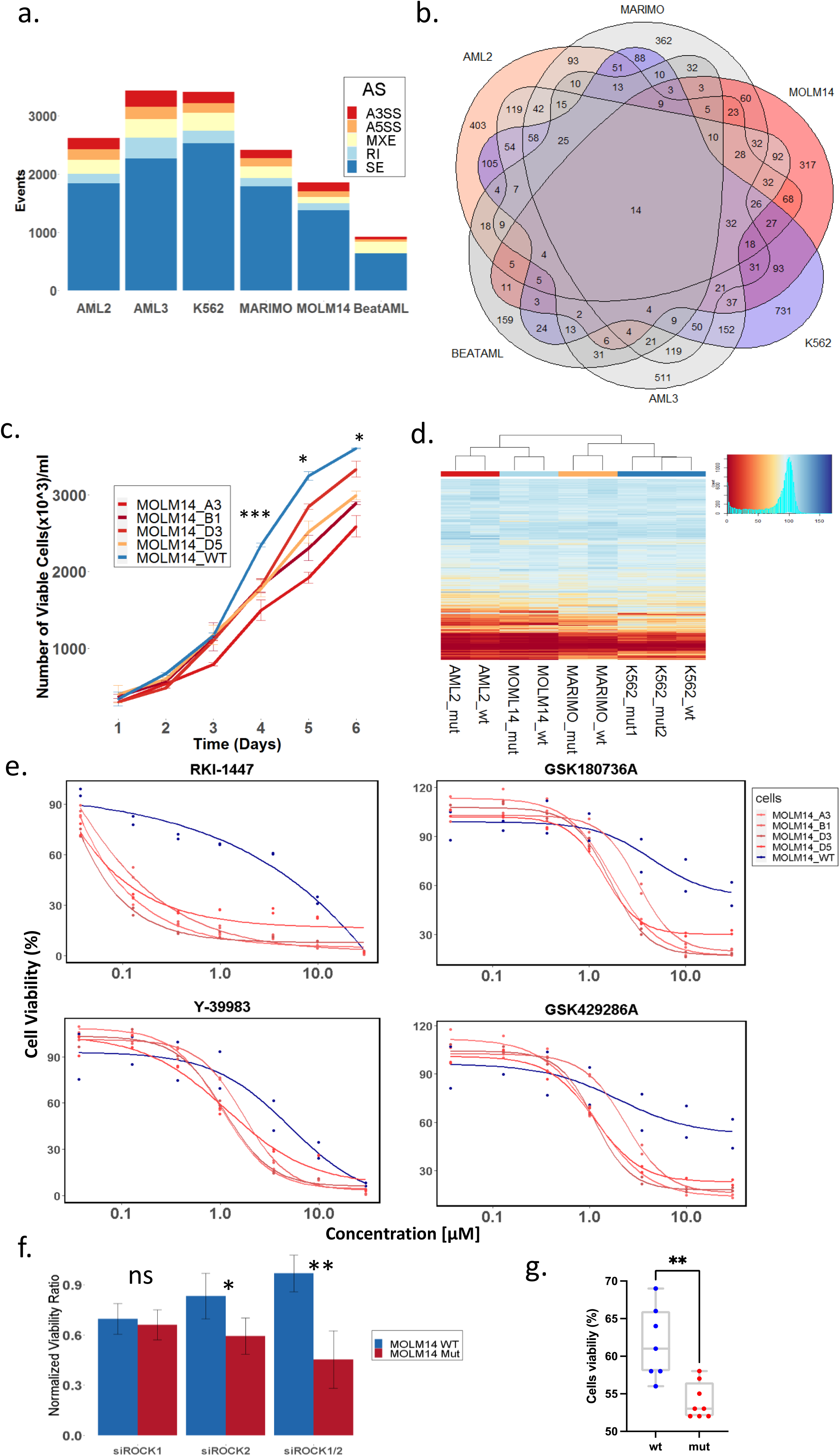
Establishment of *SRSF2* Mut models and drug screen. **a**, increased number of aberrantly spliced events was noted in all *SRSF2* Mut lines (AML2, AML3, K562, MARIMO, MOLM14), and in BeatAML *SRSF2* Mut samples (N=36) in comparison to *SRSF2* WT samples. Alternative 3′ splice sites (A3SS), alternative 5′ splice sites (A5SS), mutually exclusive exons (MXE), Retained intron (RI) and skipped exons (SE). **b**, Venn gram of aberrant exon usage (SE events) identified significant overlap (p<10e6 hypergeometric test see methods) of 14 genes with aberrant splicing in all *SRSF2* Mut samples. **c**, MOLM14 *SRSF2* Mut cells showed consistently lower growth rates than WT. The slower growth rate was significant after the fourth day in culture. **d**, Heatmap of cell viability of different isogenic *SRSF2* Mut cell lines and their WT controls with 3988 compounds (10uM). Blue represents high viability, and red represent low viability. **e**, IC-50 curve of WT, and four SRSF2 Mut MOLM14 cell lines with four different ROCKi compounds (RKI-1447, GSK180736A, Y-39983 and GSK429286A) at concentration of 0.037uM, 0.129uM, 0.369uM, 0.997uM, 29.900uM. **f**, Cell counts 48hr after transfection with siRNA targeting ROCK1 (250nM) or ROCK2 (250nM) on MOLM14 WT and *SRSF2* Mut cells. Cell counts are normalized to non-targeting siRNA controls from each group. A significant reduction in cell viability was noted after knocking down ROCK2 or knocking down both ROCK1 and ROCK2 (250nM). **g**, Isogenic *SRSF2* WT and P95L iPSC-derived HSPCs were cultured with DMSO or RKI-1447 (3μM) for 48 hours and viability was measured with CellTiter-Glo, normalized to DMSO-treated in each group. T-test, *P<0.05; **P<0.005; ***P<0.0005.

Another known phenotypic consequence of *SRSF2* mutations is a decrease in growth rate of mutant versus WT cells^10^. To validate these results, we studied the growth kinetics of our isogenic cell lines and indeed observed a slower growing rate in OCI-AML2, OCI-AML3 and MOLM14 mutated lines compared to their isogenic WT controls (**Fig.1c, Supplementary Fig. 1**).

Following these validations, we performed an HTDS of 3988 chemical compounds in a single dose (10μM), to the majority of which both *SRSF2* Mut and WT cell lines responded similarly (**Fig. 1d, Supplementary Table 4**). In addition, OCI-AML2 and MOLM14 shared a similar drug response. Following our primary screening, we identified compounds that were more cytotoxic to mutant versus WT cells (defined as a difference in viability greater than 10%, in at least two lines). A seven-dose IC50 assay was performed on three isogenic cell lines, using the selected compounds (**N=44 compounds, Supplementary Table 5**). Four of these 44 compounds were ROCKi. The IC50 analysis revealed that MOLM14 *SRSF2* Mut lines were sensitive to four different ROCKi, including GSK429286A, GSK180736A, Y-39983 and specifically to RKI-1447. OCI-AML2 *SRSF2* Mut lines were sensitive to three of them (RKI-1447, GSK429286A, GSK180736A) (**Fig. 1e, Supplementary Fig. 2, Supplementary Table 6**). RKI-1447, our leading compound (Fig. 1e), inhibits both *ROCK1* and *ROCK2*^15^. In order to exclude off-target activity and to clarify the role of *ROCK1* and *ROCK2*, we silenced both genes using a siRNA pool. Cell viability was assessed 48 hours after nucleofection. A significantly lower cell count was observed compared to WT controls after silencing *ROCK2,* or, both *ROCK1* and *ROCK2* in MOLM14 Mut cells, (**Fig. 1f**). To further validate the efficacy of RKI-1447 against human preL-HSPCs, we tested the drug on HSPCs derived through in vitro differentiation from an iPSC line carrying *SRSF2* P95L and isogenic control WT iPSC-HSPCs^7,16^. *SRSF2* Mut cells were more sensitive to RKI-1447(**Fig.1g**). Overall, by performing HTDS on our four different hematopoietic isogenic *SRSF2* mut cell lines and an iPSC line carrying *SRSF2* P95L, we identified that some *SRSF2* mut cells are sensitive to ROCKi *in vitro*.

However, to validate the relevance of these results in a more clinically relevant setting we tested the effect of RKI-1447 on primary human cells carrying the *SRSF2* Mut.

### *In vivo* Efficacy and Safety of RKI-1447 in primary human *SRSF2* Mut/WT samples

To study the efficacy of RKI-1447 on human samples, we first calibrated an *in vivo* model by injecting MOLM14 *SRSF2* Mut cells into NSG mice (**Supplementary Fig. 3a**). Once we identified a non-toxic and efficacious dosage, we conducted six AML patient-derived xenograft (PDX) experiments, with *SRSF2* Mut samples (**Supplementary Table 7,8**). Following three weeks of treatment, the engraftment of human cells in mice bone marrow was evaluated. We isolated human CD45+ cells from the total bone marrow (BM) and performed targeted sequencing of the *SRSF2* P95 region to confirm that the treatment targeted *SRSF2* Mut cells (**Supplementary Fig. 4**). In four out of the six primary AML samples, RKI-1447 significantly reduced engraftment compared to the untreated group (**Fig. 2a**) and inhibited the engraftment of both leukemic blasts and pre leukemic-HSPCs(preL-HSPCS) (**Supplementary Fig. 5,6, Supplementary Table 8**). At the same time, RKI-1447 had no significant effect on the engraftment of healthy CD34+ cells in to NSG mice (**Fig. 2b**).

**Figure 2:**
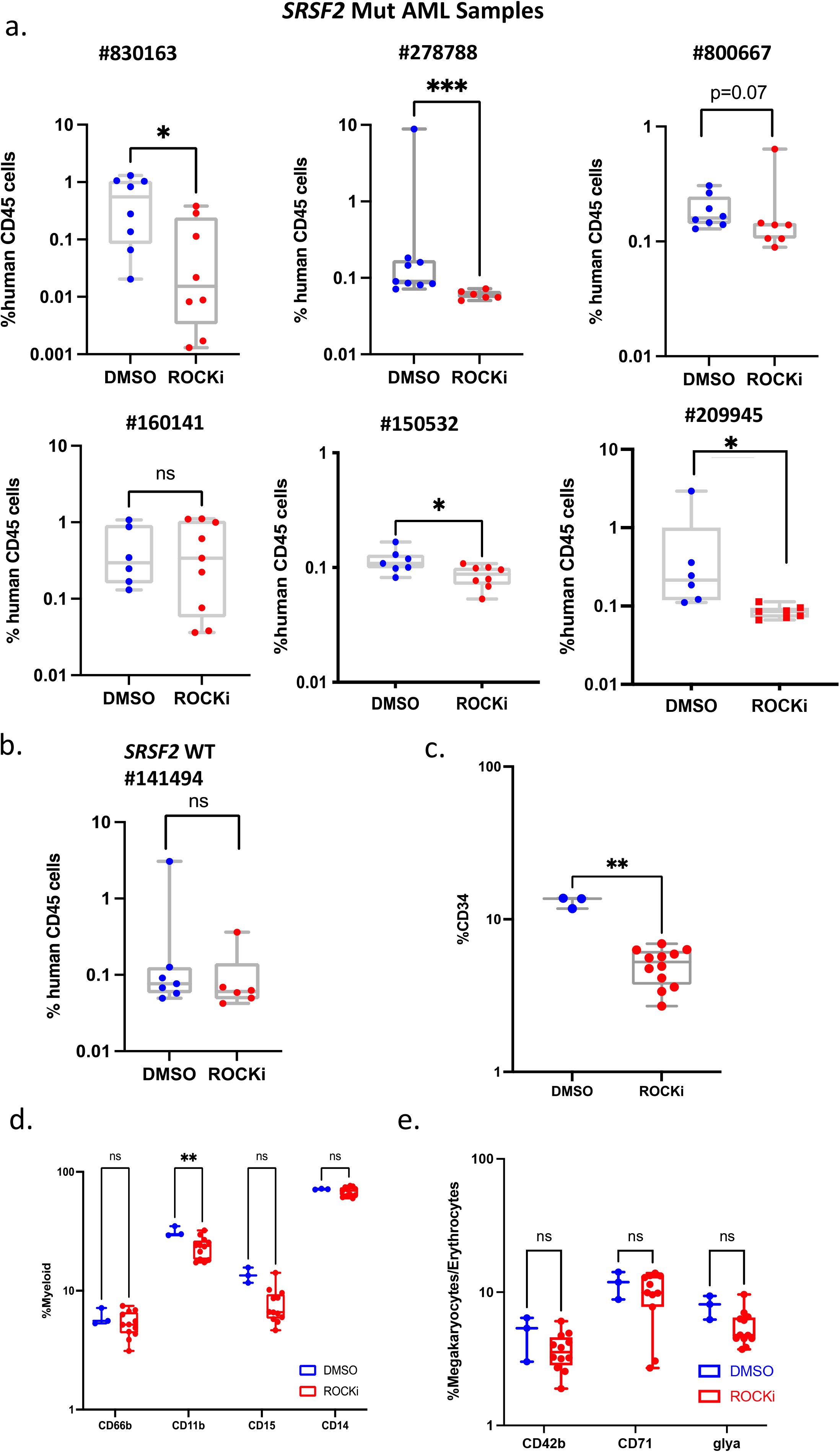
*in vivo* efficacy and safety of RKI-1447. **a**, NSG mice were treated with RKI-1447 (50mg/kg) for 3 weeks after transplantation of CD3 depleted human cells from six patients with *SRSF2* mut acute myeloid leukemia (AML). The percentage of human CD45+ (hCD45) cells in engrafted murine bone marrow with/without RKI-1447 treatment is shown after staining for CD45, CD33, CD3 and CD19 to determine myeloid or lymphoid lineage cells and analyzed by flow cytometry. **b**, NSG mice were treated with RKI-1447(50mg/kg) for 3 weeks after transplantation of CD34+ cells from healthy individual (SRSF2 WT). The percentage of human CD45+ (hCD45) cells in engrafted murine bone marrow with/without RKI-1447 treatment is shown after staining for CD45, CD33, CD3 and CD19 to determine myeloid or lymphoid lineage cells and analyzed by flow cytometry. **c**, Percentage of CD34+ cells by *ex vivo* culturing of healthy CD34+ cells with exposure to RKI-1447 at different concentration (3μM, 1μM, 0.33μM, 0.11μM, 0.04μM, 0.01μM). **d-e**, *ex vivo* differentiation of healthy CD34+ cells towards myeloid (**d**), megakariod and erythroid cells (**e**) with different cytokines (see method) after exposure to RKI-1447 at different concentration (3μM, 1μM, 0.33μM, 0.11μM, 0.04μM, 0.01μM). Myeloid lineage is detected with CD66b, CD11b, CD15, CD14; Megakaryocytes are detected with CD42b; Erythrocytes are detected with CD71 and GlyA. Mann−Whitney U test with FDR correction for multiple hypothesis testing, *P<0.05; **P<0.005; ***P<0.0005.

Once we established that RKI-1447 was active against human samples carrying *SRSF2* Mut we aimed at testing its toxicity against different human tissues *in vitro*. We isolated human CD34+ HSPCs, seeded them in 384-well plates and induced differentiation toward myeloid, erythroid and megakaryoid lineages. After seven days, RKI-1447 was added, cells were further incubated for 48 hours, and cell viability was measured. RKI-1447 demonstrated limited toxicity to the three lineages treated (**Supplementary Fig. 7a**). Flow cytometry analysis of colonies after RKI-1447 treatment demonstrated that there were less CD34+ and CD11b+ cells (**Fig. 2 c, d**), but differentiation was normal towards erythrocytes and megakaryocytes (**Fig.2 e**), and other myeloid populations.

Liver metabolism assay revealed that RKI-1447 is metabolized in the liver with a half-life of 11.42 minutes (**Supplementary Fig. 7b**) and hepatotoxicity was comparable to Tamoxifen effect (**Supplementary Table 9**). We also evaluated the potential cardiotoxicity with the hERG assay (a predictor assay based on a compound binding to hERG channel, which can lead to cardiotoxicity). Compared with Cisapride, a known cardio-toxic compound, RKI-1447 was predicting to have very minor cardiotoxicity (**Supplementary Fig. 8a**). To test whether the compound induced differentiation of our MOLM14 *SRSF2* mut model, we performed flow cytometry before and after 24 hours’ exposure to RKI-1447 (0.5μM). Examination of different differentiated blood cell markers indicated that there was no difference in differentiation between treated and non-treated cells (**Supplementary Fig. 8b-e**). Neutrophil differentiation related genes *AZU1*, *CTSG*, *ELANE* and *MPO* were not differentially expressed after exposure to RKI-1447 in *SRSF2* Mut MOLM14 cells (**Supplementary Fig. 8f**). The expression of *CTSG*, *ELANE* and *MPO* were even down regulated in *SRSF2* WT after exposure to RKI-1447 (**Supplementary Fig. 8g**). Overall, we validated the efficacy and toxicity of RKI-1447 in *SRSF2* mutated AML samples using xenograft models and toxicity assays. As we established the efficacy of RKI-1447 next we aimed at understanding its mechanisms of action.

### ROCK inhibitors cause Mitotic Stress

To study the molecular and cellular mechanisms underlying the response of SRSF2 Mut to ROCK inhibition. We exposed MOLM14, OCI-AML2 and MARIMO isogenic lines to RKI-1447 and measured the proteomic and RNA-seq profiles before exposure to the drug and two or eight hours later. Unbiased analysis of the proteomic data demonstrated a clear separation of protein expression between Mut and WT cells at all time points of all cell lines (**Fig. 3a, Supplementary Fig. 9**). In MOLM14 we observed a time dependent proteome expression change in both Mut and WT (**Fig. 3a**). These results suggest that RKI-1447 has a clear effect on the proteome of the Mut and WT cells; however, their baseline differences are maintained. Furthermore, gene-set enrichment analysis of the protein expression between *SRSF2* Mut before and after exposure to RKI-1447 identified cell cycle and G2M checkpoint pathways as significantly upregulated among cells eight hours after treatment (**Fig. 3b, Supplementary Fig. 10**). The upregulation of mitosis related genes in the MOLM14 *SRSF2* Mut after RKI-1447 (**Fig. 3c**) is a sign of mitotic arrest. In accordance with these findings, treatment with RKI-1447 resulted in a higher percentage of cells in G2/M (**Fig.3d**). Similar results were noted for MOLM14 *SRSF2* WT cells (**Supplementary Fig. 11**). Suggesting that RKI-1447 can cause similar effects in both *SRSF2* Mut and WT MOLM14, but for some reason *SRSF2* Mut cells are more sensitive to the mitotic stress. Rho GTPases are important regulators of mitosis and cytokinesis in mammalian cells through their ability to modulate microtubule dynamics and actin-myosin cytoskeleton^17,18^. Accordingly, we stained our *SRSF2* Mut and WT cell lines with anti-α-tubulin and DAPI. We observed significantly more abnormal multipolar mitotic spindles in *SRSF2* Mut cells compared with WT after exposure to RKI-1447 (**Fig. 3e, f**). We also observed an increase in multi-nuclear cells (**Fig. 3f**). These findings suggest that ROCKi induces abnormal mitosis correlated with abnormal microtubule organization, supporting our proteomic and cell cycle observations. To further explore the nature of the differential effect of RKI-1447 on the WT and *SRSF2* Mut cells, a comprehensive 3-dimensional characterization of the cytoskeleton and the nucleus in these cells was conducted.

**Figure 3:**
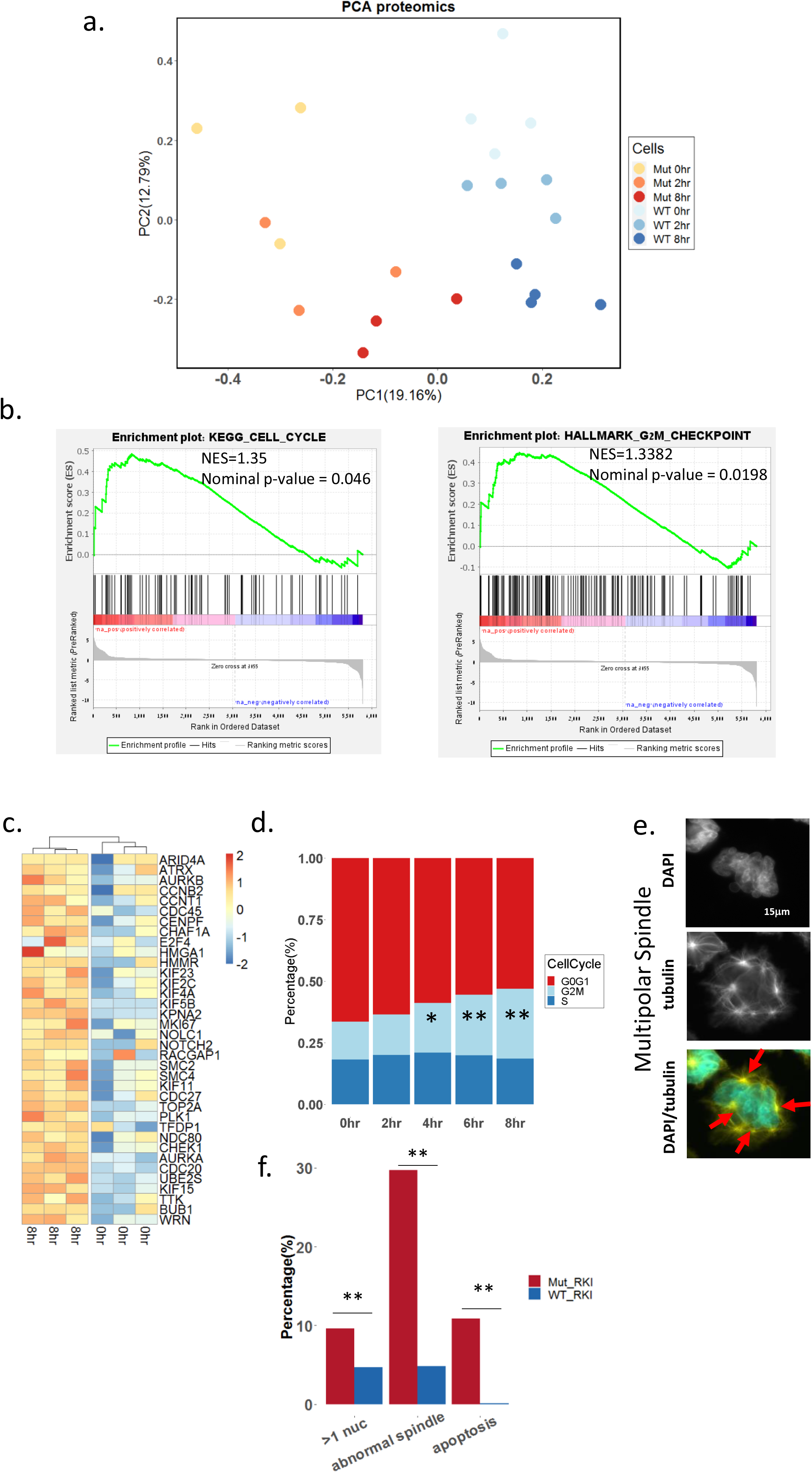
RKI-1447 affect cell cycle and cause mitotic stress of *SRSF2* Mut. **a**, Principal component analysis plot of protein expression of MOLM14 WT and *SRSF2* Mut cells before (0hr) and after 2 (2h) and 8 (8h) hours of exposure to RKI-1447 (0.5μM). **b**, A GSEA analysis of protein expression pre-ranked based on the LOG2 fold change between time before exposure (0h) and eight hours after exposure to RKI-1447 (0.5μM) on MOLM14 *SRSF2* Mut cells. Significant enriched pathways from the GSEA analysis included the Cell Cycle and G2M checkpoint pathways. **c**, Heatmap of core enrichment genes for gene set G2M checkpoint between MOLM14 *SRSF2* Mut before and after exposure to RKI-1447 (0.5μM) 8hr. **d**, Cell cycle of MOLM14 *SRSF2* Mut before and after 2hr, 4hr, 6hr, and 8hr exposure to RKI-1447(0.5μM). Different cell cycles are indicated with different colors in the plot. Significant increased G2M is observed after 4hr, 6hr and 8hr exposure to RKI-1447. T-test, *P<0.05; **P<0.005; ***P<0.0005. **e**, Representative image of abnormal spindles in MOLM14 *SRSF2* Mut cells after 24 hours exposure to RKI-1447 (0. 5μM). Cells are stained with DAPI and α-tubulin. **f**. Percentage of cells with multi nucleis, abnormal spindles and apoptosis in MOLM14 *SRSF2* Mut and WT after 24hr exposure to RKI-1447 (0.5μM). Pearson’s chi-square, *P<0.05; **P<0.005; ***P<0.0005.

### Nuclear Deformations and Cytoskeleton Disruption

As the cytoskeleton is one of the major contributors to nuclear shape^19^ and we noticed abnormal microtubule organization among *SRSF2* Mut cells after exposure to RKI-1447, we next examined the general effects of the mutation and the RKI-1447 on the cells’ structure. Three-dimensional confocal imaging of WT and mutant cells, either untreated or treated with RKI-1447, were performed and revealed major differences in general nuclear morphology both before RKI-1447 and after.

In WT cells the DAPI-labeled nuclei showed some heterogeneity, ranging from round to cashew-nut morphology (**Supplementary Fig. 12a**). Mutant cells displayed higher level of heterogeneity, with larger nuclei, displaying multiple indentations (**Supplementary Fig. 12a**). Rendering the WT and mutant nuclei, followed by quantitative morphometric analysis, indicated that the untreated mutant cells displayed a significant larger mean surface area (**Supplementary Fig. 12b**) and larger mean volume (**Fig. 4a, Supplementary table 10**) than the untreated WT cells. Upon treatment with RKI-1447 of MOLM14 WT cells, major changes in the overall shape of the nucleus occurred, manifested by nuclear lobulation (**Supplementary Fig. 12a**). Treatment of the mutant cells with RKI-1447, increased the nuclear indentations and induced extensive nuclear lobulation (**Supplementary Fig. 12a**). Upon the treatment of RKI-1447, the surface area and the volume of the MOLM14 WT cells were significantly increased and were similar to that of the Mut. These data suggest that the nuclear indentations and segmentation induced by *SRSF2* mutations at baseline are augmented by RKI-1447. To explore the nature of the structural phenotype associated with the *SRSF2* mutation, at higher resolution, we subjected the WT and mutant cells (RKI-1447-treated and un-treated), to comprehensive TEM imaging (**Fig. 4b and Supplementary Fig. 13**). WT cells displayed nuclei with variable morphology, commonly containing conspicuous indentations, shown here in longitudinal section (**Fig. 4b and Supplementary Fig.13a**). Consistent with the rendered data, treatment of WT cells with RKI-1477 resulted in increased nuclear indentations (**Figure 4b, Supplementary Fig. 13a**). Untreated *SRSF2* mutant cells displayed deep invagination, in which the cytoplasmic intrusions essentially “went all the way”, effectively producing extended regions in which nuclear membrane areas apparently interact with each other via the associated nuclear laminae (**Fig. 4b, Supplementary Fig.13b**). Addition of RKI-1447 to the mutant cells resulted in hyper-segmentation and hyper-lobulation of the nucleus (**Figure 4b, Supplementary Fig. 13b**). This nuclear morphology is reminiscent of the nuclear shape of Pelger–Huët anomaly^20^. Altogether, adding SRSF2 mutations to MOLM14 created a pseudo Pelger–Huët anomaly nuclear shape.

**Figure 4.**
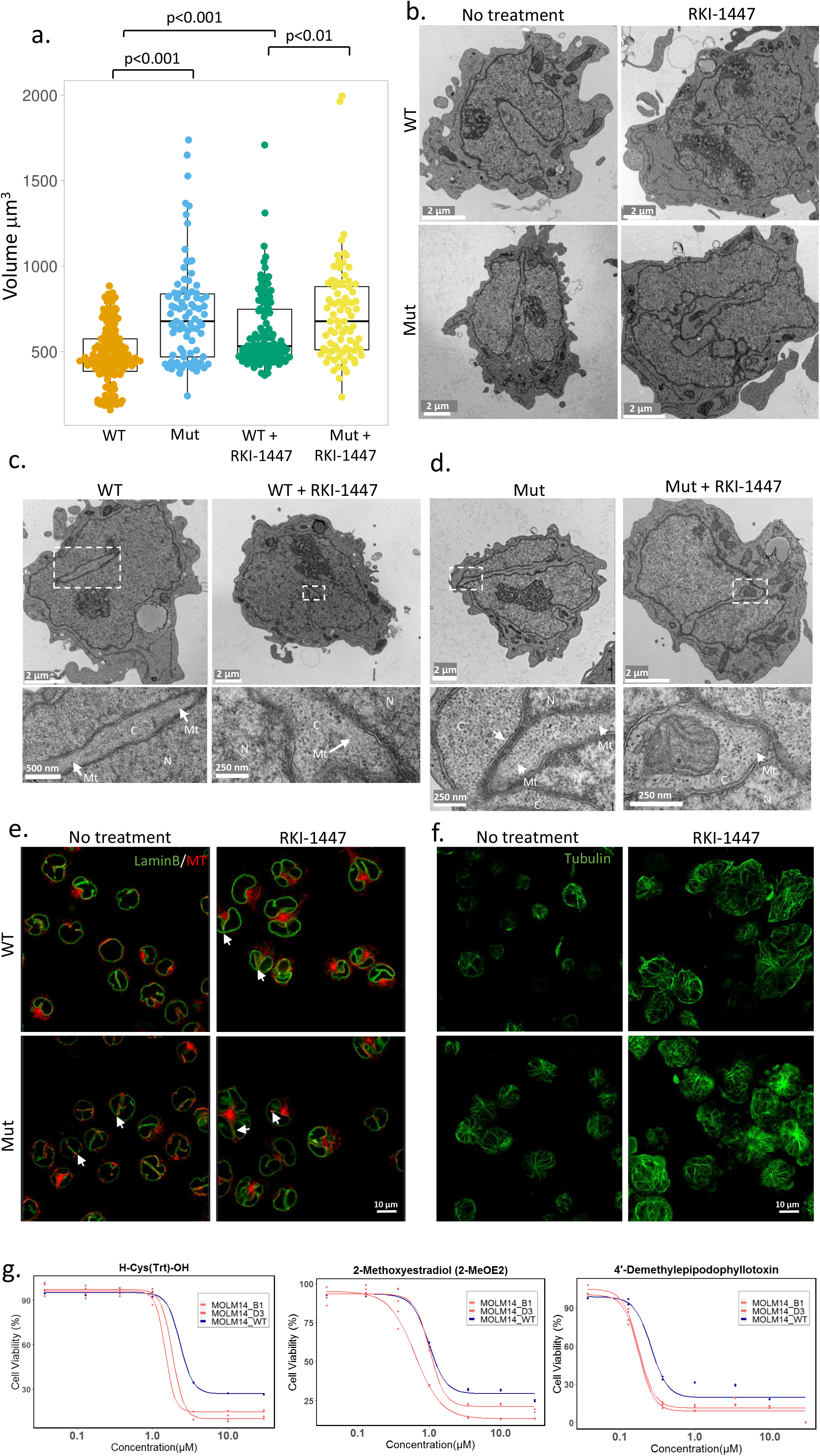
Confocal and transmission electron microscopy images of WT and *SRSF2* Mut MOLM14 cells before and after exposure to RKI-1447. WT and *SRSF2* Mut MOLM14 cells were either left untreated or treated with 0.5μM RKI-1447 for 24 hours before fixation as indicated. **a**, 3D confocal images of DAPI labeled nuclei were subjected to rendering and morphometric quantifications. Nuclear volume is presented in box and whiskers plot format. **b**, Transmission electron microscopy (TEM), showing a common presence of a deep (“half-way”) nuclear indentation (top-left image); in mutant cells, these nuclear indentations were considerably deeper, usually contacting the opposite side of the nucleus, thereby segmenting the nucleus (bottom-left image); Treatment of WT cells with RKI-1447 induced conspicuous nuclear segmentation (top-right image). Note that segments are often connected to each other by extended sheets consisting of two nuclear membranes, inter-connected by the associated nuclear laminae; “hyper-segmentation” of the nuclei of the mutant cells (bottom-right). For higher magnification view, see also **Supplementary Fig 13. c-d**, TEM images illustrate the overall morphology of *SRSF2* WT (**c**) and mutant (**d**) cells, untreated or treated with RKI-1447 (0.5μM), are shown in lower magnification (top panel) and higher magnification (bottom panel, corresponding to the white rectangles in the top panel). Nuclear regions and cytoplasmic regions are marked C and N respectively. Arrows marked by “Mt” point to microtubules that are associated with the edges of the cytoplasmic insertions into the nucleus. The arrow in the “mut” (bottom panel) points to the “interlobular sheet” consisting of the two nuclear membranes and the nuclear lamina running between them. **e**, Cells were labeled with anti-LaminB1 antibodies to outline nuclear membrane (shown in green) and anti-α-tubulin antibodies to label microtubules (shown in red). 3D volumes were acquired on the Olympus confocal microscope. Representative confocal slices corresponding to the middle plane of the cells are shown. Note, deep narrow folds of nuclear membrane that are especially prominent in mutant cells and upon RKI-1447 treatment. Red signal detected inside such invaginations points to the presence of microtubules (arrows). See also **Supplementary movie 1** and **Supplementary Fig.14. f**, Representative Images of microtubules (shown in green), in *SRSF2* WT and Mut cells, untreated or treated with RKI-1447 (0.5μM), were acquired using the Leica SP8 scanning confocal microscope. Cells were labeled with anti-α-tubulin antibodies. A series of confocal slices depicting the overall morphology of the microtubular network in these cells is shown in **Supplementary Fig.15. g**, Dose response curves of four cytoskeleton inhibitors against *SRSF2* WT and Mut MOLM14 cell lines. Viability of *SRSF2* WT and Mut MOLM14 cell lines were measured after 48 hours exposure to three microtubule modifiers: 4’-Demethylepipodophyllotoxin, H-Cys(Trt)-OH, 2-Methoxyestradiol at concentration of 0.037μM, 0.129μM, 0.369μM, 0.997μM and 29.90μM.

Interestingly, higher power magnification of the cytoplasmic intrusions into the nucleus (**Fig. 4c (WT), Supplementary Fig. 13a’**) revealed arrays of microtubules in the proximity of the indented nuclear membrane. Notably, the “leading edges” of the cytoplasmic intrusions in ROCKi-treated WT cells, contained arrays of microtubules too, similar to those found in untreated WT cells (**Fig. 4c (WT +RKI-1447), Supplementary Fig. 13a’**). High power examination of the *SRSF2* Mut cells, reveal microtubules that are attached to the tips of the cytoplasmic intrusions into the nucleus, and apparently “push on” the nuclear membrane and indents it (**Fig. 4d (Mut), Supplementary Fig. 13b’**). Section of broad cytoplasmic intrusions, that are prominent in ROCKi-treated SRSF2 Mut cells, reveal extended areas where the nucleoplasm was essentially dissected by the “invading” microtubule-rich cytoplasm (**Fig. 4d (Mut +RKI-1447), Supplementary Fig. 13b’**).

The transmission EM data provided high resolution visualization of the nuclear indentations and segmentation, yet could not provide comprehensive 3D view of the nucleus-cytoplasm interplay and topology in these cells. To achieve such information, we immunolabeled WT and *SRSF2* mut MOLM14 cells (RKI-1447-treated and untreated) with tubulin and laminB1, which delineates the nuclear membrane.

### Abnormal microtubule system and the sensitivity to microtubule modifiers in *SRSF2* Mut

The LaminB1 and tubulin staining reinforce the TEM data, directly demonstrating that that the *SRSF2* Mut and RKI-1447 treatment strongly enhance the indentation of the nuclear membrane by modulating microtubule network (**Fig. 4 e, f, Supplementary Fig. 14,15 and Supplementary Movie 1**). Based on the imaging studies, we suspected that *SRSF2* mut cells would also be sensitive to other microtubule modifiers that would break the balance of the microtubule system. We tested 34 compounds, which can modulate/bind either tubulin, myosin or actin. We identified four more compounds, which were significantly more toxic to *SRSF2* Mut versus WT (**Fig. 4g, Supplementary Fig. 16**). Three of these compounds are microtubule inhibitors: 4’-Demethylepipodophyllotoxin (a potent inhibitor of microtubule assembly)^21^, H-Cys(Trt)-OH (a specific inhibitor of Eg5 that inhibits Eg5-driven microtubule sliding velocity in a reversible fashion)^22^, 2-Methoxyestradiol (an inhibitor of polymerization of tubulin)^23^, and the other one is myosin II inhibitor, Blebbistatin ^24^ (**Supplementary Fig. 16**).

### Nuclear F-Actin is increased in *SRSF2* Mut and modified by RKI-1447

As ROCK also controls actin polymerization and stabilization (mostly through its effects on acto-myosin II contractility), we have labeled WT and mutant cells (untreated and RKI-1447 treated) with DAPI, phalloidin. An increase of nuclear F-actin was observed in *SRSF2* mut cells, compared to *SRSF2* WT at baseline and without treatment. (**Supplementary Fig. 17a**). Following addition to RKI-1447, the ratio of nuclear actin/cytoplasm actin was increased from 0.52 to 0.68 in WT cells, while it was decreased from 0.86 to 0.67 in SRSF2 mut cells (**Supplementary Fig. 17b**). These findings expose yet another cytoskeleton system which is different between *SRSF2* Mut and WT cells.

Taken together, the discovery of *SRSF2* Mut cells sensitivity to RKI-1447 exposed novel phenotypes of the *SRSF2* Mut cells. These phenotypes (abnormal cytoskeleton organization of both microtubules and F-Actin) are augmented by RKI-1447 treatment and thus explain why *SRSF2* Mut cells are sensitive to RKI-1447.

## Discussion

In the current study, we aimed at designing new models for HTDS against *SRSF2* Mut cells (**Fig. 1**) with the global aim of identifying novel safe and efficacious targeted therapies against *SRSF2* Mut in humans (**Fig. 2**). We discovered that ROCKi were effective against *SRSF2* Mut cells lines and that the leading ROCKi was effective in targeting human AML cells carrying *SRSF2* mutations in a xenograft model (**Fig. 2**). The study of synthetic lethality can help not just in identifying novel targets but also in understanding the mechanisms underlying *SRSF2* mutations (**Fig. 3**). Although *SRSF2* Mut has generally been assumed to cause disease by inducing specific splicing defects, here we provide evidence that they also cause cytoskeletal and nuclear morphological abnormalities (**Fig. 4**) which are augmented by the ROCKi to a level were the cells develop such abnormal nuclei morphology, and spindle irregularity, which are incompatible with cell survival (**Fig. 3,4**).

Our study has both clinical and molecular mechanistic implications. From the clinical point of view, we identified new targets against *SRSF2* Mut cells that have been validated on primary human AML samples. Our comprehensive approach was fruitful as we extended the diversity of cell lines, we used for the drug screen. While our model system exposed sensitivity to several different ROCKi, they were only effective in two out of four of the cell lines tested, and in four out of six of the primary AML samples. While RKI-1447 is not optimized for a particular clinical setting, we were able to demonstrate its safety (**Fig. 2**), therefore we propose it should be considered as a clinical trial candidate.

From the mechanistic point of view, we discovered that RKI-1447 cause mitotic arrest and abnormal spindles in *SRSF2* Mut cells, which, if left unresolved lead to cell death (**Fig. 3**). In order to understand why *SRSF2* Mut cells were more sensitive to RKI-1447 induced mitotic stress, we extended our study to the cytoskeleton and nuclear morphology which are tightly regulated by the Rho-ROCK pathway^25^. Towards that end we stained F-actin sub-cellular distribution (which is known to be regulated by ROCK)^26^, and found increased nuclear F-actin in *SRSF2* Mut cells, which is correlated with DNA damage^27^. Previous studies have suggested that *SRSF2* Mut have cell cycle abnormalities, and that they might be sensitive to cell cycle modulation due to DNA replication stress and R-loop formation^9,28,29^.

Next, we performed quantitative analysis of nuclear morphology, which exposed significantly higher nuclear volume and cell surface area of *SRSF2* Mut cells compared to WT cells at baseline (before treatment) (**Fig.4a, Supplementary Fig. 12a, b**). To gain better understanding of the cytoskeletal structures that can induce such differences, we have used TEM, which revealed a segmented pseudo Pelger–Huët anomaly (PHA) nuclei. Pelger–Huët anomaly is a blood laminopathy associated with mutations in the lamin B receptor (LBR) gene^30^, and the characteristic of which is bi-lobed nuclei of leukocytes with symmetrical “dumbbell” or “pince-nez” appearance are connected by a thin filament of chromatin^20^. Acquired pseudo-PHA is seen also in the MPN, MDS, AML, infections and medications. While pseudo PHA is a phenotype of mature neutrophils in the current study, we provide evidence that the introduction of *SRSF2* mutations to AML blasts (myeloid progenitors) can cause the pseudo-PHA phenotype. To investigate why *SRSF2* Mut cause the structural changes we observed, we looked into the cytoplasmic intrusions that segment the nuclei, and the structures that connect the lobes of the nuclei by high power TEM. This examination revealed enrichment of microtubule bundles inside the nuclear indentations (**Fig. 4 c,d**). This result was further supported by LMNB1 and microtubules staining (**Fig.4 e,f, Supplementary movie1**). While our findings expose abnormal nuclear shape and hyperactive microtubules, it remains elusive why *SRSF2* Mut and possibly other splicing mutations modify the nucleoskeleton, at the molecular level. We believe that this is one of the first reports providing evidence that the microtubule network is more prominent in *SRSF2* Mut, especially after ROCKi treatment. More importantly, these microtubules were located at the tip of the cytoplasmic intrusions suggesting that they take an active role in the nuclear segmentation process.

Based on our results, we suggest that the hyperactive microtubule system might be the cause of pseudo-PHA. Future studies are needed to explore the efficacy of ROCKi in MDS and AML with dysplastic changes and the role of microtubules in MDS and other SMMs. In this regard, a study with siRNA screening in AML samples identified ROCK as a target in AML with efficacy in 20% of primary AML samples ^31^. Our results suggest that modulating the cytoskeleton by ROCKi and possibly other microtubule modifiers (**Fig. 4g**) can take advantage of the abnormal cytoskeleton induced by *SRSF2* Mut, and open new ways to treat AML/MDS and pre-leukemia.

## Acknowledgements

This research was supported by the National Natural Science Foundation of China (81861148029, 81730006, 81890990), National Key R&D Program of China (2020YFE0203000, 2021YFA1100900), CAMS Innovation Fund for Medical Sciences (CIFMS) (2021-I2M-1-040), CAMS Fundamental Research Funds for Central Research Institutes (3332021093), the EU horizon 2020 grant project MAMLE ID: 714731, LLS and rising tide foundation Grant ID: RTF6005-19, ISF-NSFC 2427/18, ISF-IPMP-Israel Precision Medicine Program 3165/19, ISF 1123/21, BIRAX 713023, the Ernest and Bonnie Beutler Research Program of Excellence in Genomic Medicine, awarded to LIS. LIS is an incumbent of the Ruth and Louis Leland career development chair. N.K. is an incumbent of the Applebaum Foundation Research Fellow Chair. This research was also supported by the Sagol Institute for Longevity Research, the Barry and Eleanore Reznik Family Cancer Research Fund, Steven B. Rubenstein Research Fund for Leukemia and Other Blood Disorders, the Rising Tide Foundation and the Applebaum Foundation. Work in the Papapetrou lab was supported by US National Institutes of Health (NIH) grants R01HL137219 and R01CA225231, the Leukemia and Lymphoma Society and the Edward P Evans Foundation.

Electron microscopy studies were conducted at the Irving and Cherna Moskowitz Center for Nano and Bio-Nano Imaging at the Weizmann Institute of Science.

Some of the images in this paper were acquired at the Advanced Optical Imaging Unit, de Picciotto-Lesser Cell Observatory unit at the Moross Integrated Cancer Center Life Science Core Facilities, Weizmann Institute of Science.

The SGC library was supplied by the Structural Genomics Consortium under an Open Science Trust Agreement: https://www.thesgc.org/click-trust.

## Author contribution

M.S., L.I.S., T.F. and B.G. designed the study and wrote the manuscript. M.S. performed the experiment and analyzed the data. M.S. and N.K. performed in vivo experiment and analyze the data. T.F. generated the SRSF2 isogenic cell lines and performed the experiment. I.G. and M.S. prepared the samples for imaging. M.B.H. and N.D. performed the imaging and M.B.H. analyzed the data. H.B. and A.P. performed the drug screen assay. M.O. and E.P.P. designed and performed the iPSCs experiment. M.M. contributed clinical samples. Y.M. helped with amplicon sequencing and CRISPR. N.C. help with the data analyze. T.C. help with the data interpretation. L.I.S. supervised the study.

