## Supplementary Figs for "Rock inhibitors target *SRSF2* leukemia by disrupting cell mitosis and nuclear morphology"

Supplementary Fig. 1

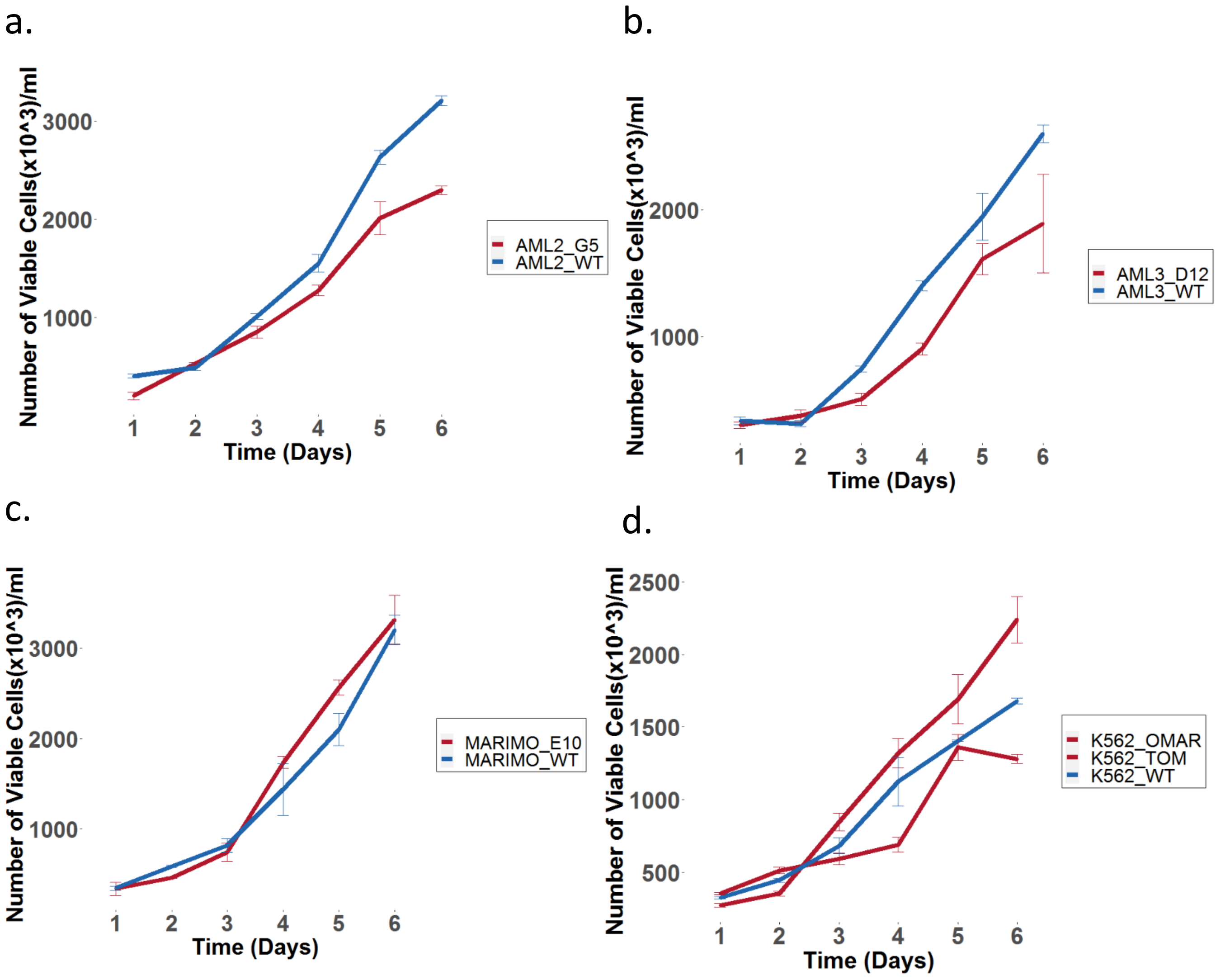

### Supplementary Fig. 2

#### OCI-AML2

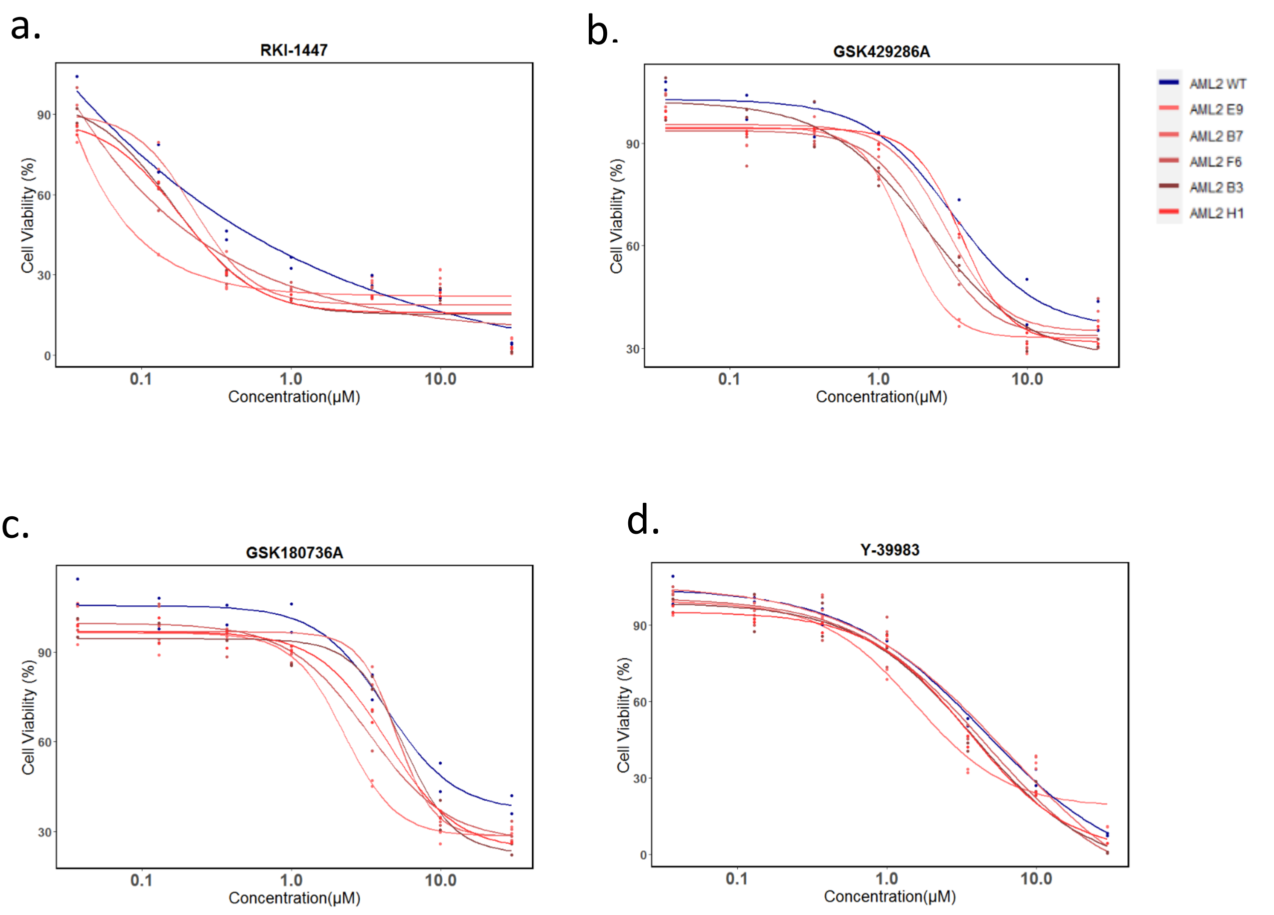


#### MARIMO

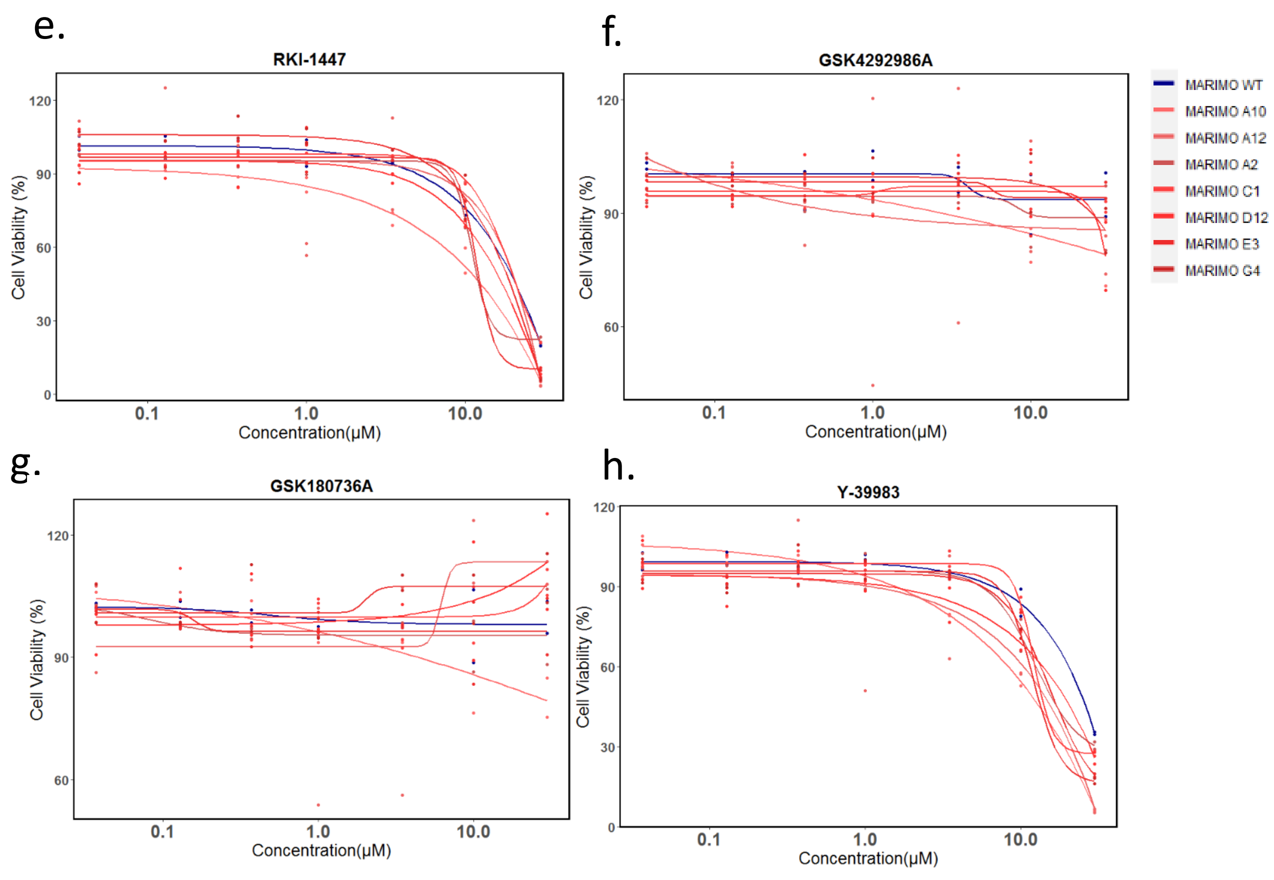

Supplementary Fig. 3

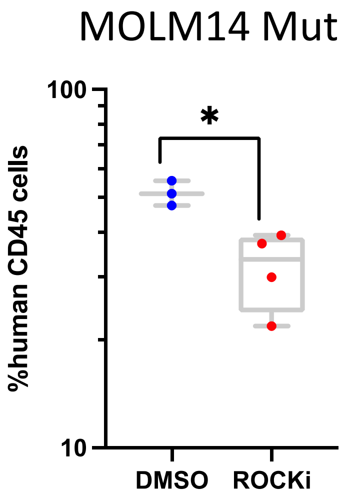

Supplementary Fig. 4

| Mouse No. | VAF | AAChange.refGene |
|---|---|---|
| 150532_DMSO_6 | 0.215 | c.C284A:p.P95H |
| 150532_DMSO_7 | 0.028 | c.C284A:p.P95H |
| 150532_RKI_11 | 0.153 | c.C284A:p.P95H |
| 150532_RKI_13 | 0.051 | c.C284A:p.P95H |
| 209945_DMSO_16 | 0.481 | c.C284G:p.P95R |
| 209945_DMSO_17 | 0.284 | c.C284G:p.P95R |
| 209945_DMSO_19 | 0.293 | c.C284G:p.P95R |
| 209945_DMSO_20 | 0.488 | c.C284G:p.P95R |
| 209945_RKI_23 | 0.419 | c.C284G:p.P95R |
| 209945_RKI_27 | 0.46 | c.C284G:p.P95R |
| 209945_RKI_28 | 0.49 | c.C284G:p.P95R |
| 160141_DMSO_2 | 0.069 | c.C284A:p.P95H |
| 160141_DMSO_3 | 0.012 | c.C284A:p.P95H |
| 160141_DMSO_4 | 0.033 | c.C284A:p.P95H |
| 160141_RKI_7 | 0.05 | c.C284A:p.P95H |
| 160141_RKI_8 | 0.035 | c.C284A:p.P95H |
| 160141_RKI_9 | 0.021 | c.C284A:p.P95H |
| 160141_RKI_11 | 0.011 | c.C284A:p.P95H |
| 160141_RKI_12 | 0.043 | c.C284A:p.P95H |
| 160141_RKI_13 | 0.009605 | c.C284A:p.P95H |
| 160141_RKI_14 | 0.073 | c.C284A:p.P95H |
| 800667_DMSO_1 | 0.478 | c.C284A:p.P95H |
| 800667_DMSO_2 | 0.515 | c.C284A:p.P95H |
| 800667_DMSO_3 | 0.445 | c.C284A:p.P95H |
| 800667_DMSO_4 | 0.402 | c.C284A:p.P95H |
| 800667_DMSO_5 | 0.224 | c.C284A:p.P95H |
| 800667_DMSO_6 | 0.14 | c.C284A:p.P95H |
| 800667_DMSO_7 | 0.471 | c.C284A:p.P95H |
| 800667_DMSO_8 | 0.202 | c.C284A:p.P95H |
| 800667_RKI_9 | 0.334 | c.C284A:p.P95H |
| 800667_RKI_10 | 0.519 | c.C284A:p.P95H |
| 800667_RKI_11 | 0.457 | c.C284A:p.P95H |
| 800667_RKI_12 | 0.5 | c.C284A:p.P95H |
| 800667_RKI_13 | 0.124 | c.C284A:p.P95H |
| 800667_RKI_14 | 0.501 | c.C284A:p.P95H |
| 800667_RKI_15 | 0.599 | c.C284A:p.P95H |
| 278788_DMSO_14 | 0.013 | c.C284A:p.P95H |
| 278788_DMSO_17 | 0.038 | c.C284A:p.P95H |
| 278788_DMSO_20 | 0.028 | c.C284A:p.P95H |
| 278788_DMSO_21 | 0.02 | c.C284A:p.P95H |
| 278788_RKI_24 | 0.03 | c.C284A:p.P95H |
| 278788_RKI_27 | 0.042 | c.C284A:p.P95H |
| 830163_DMSO_6 | 0.0706 | c.C284G:p.P95R |
| 830163_RKI_12 | 0.0284 | c.C284G:p.P95R |

Supplementary Fig. 5

a

.

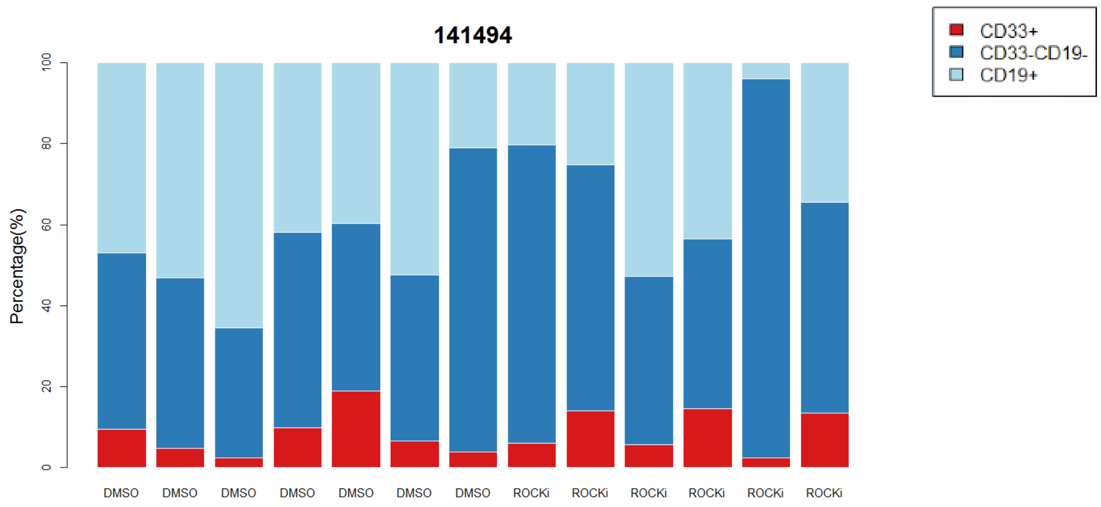


b.

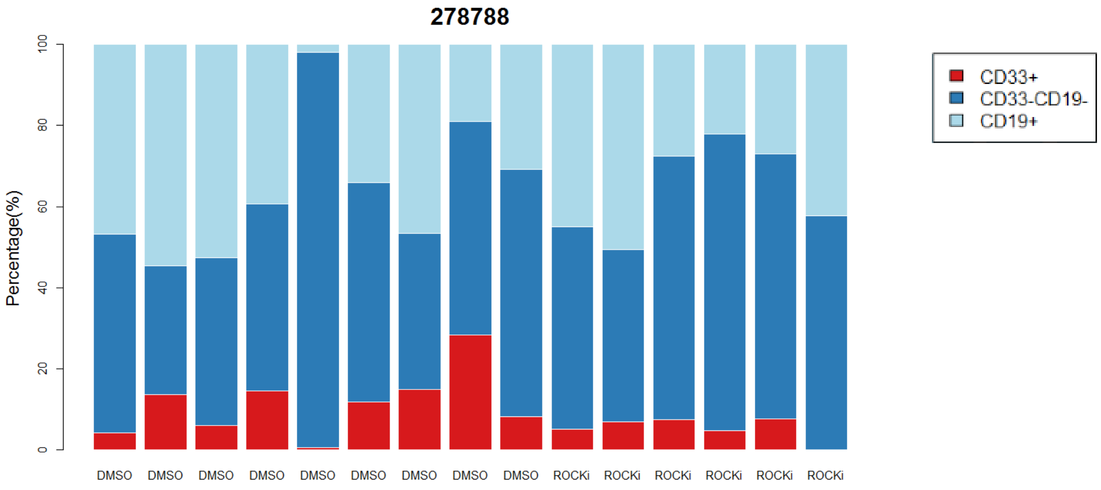


c.

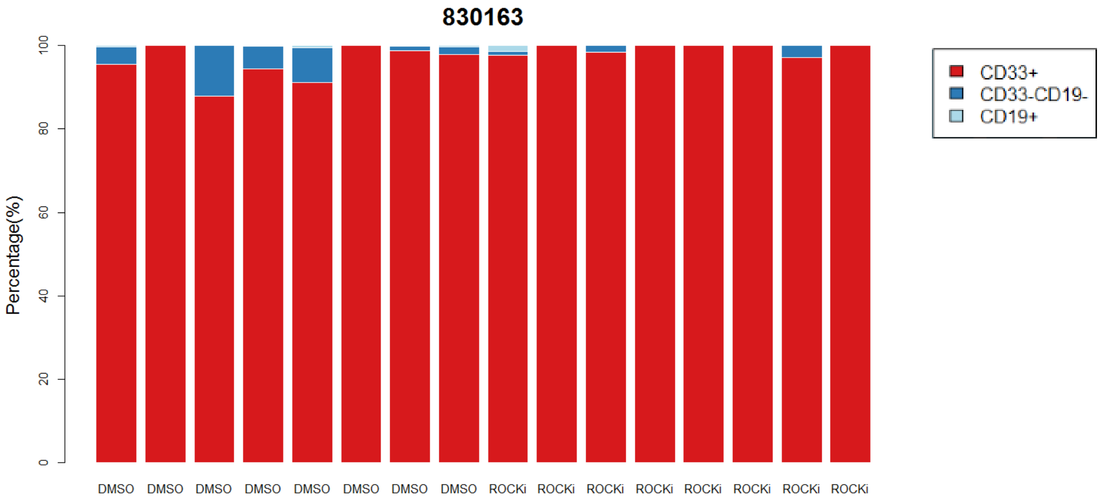


d.

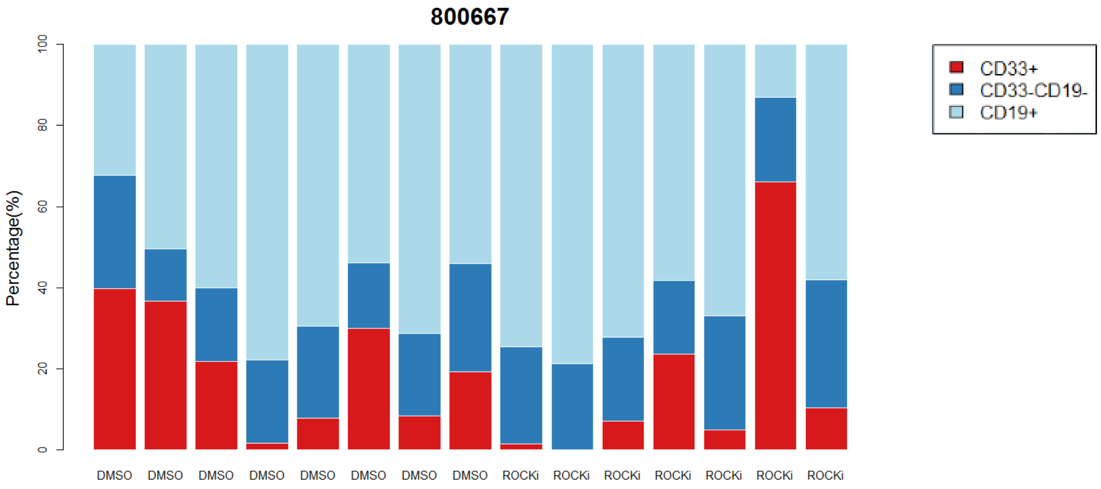

Supplementary Fig. 6

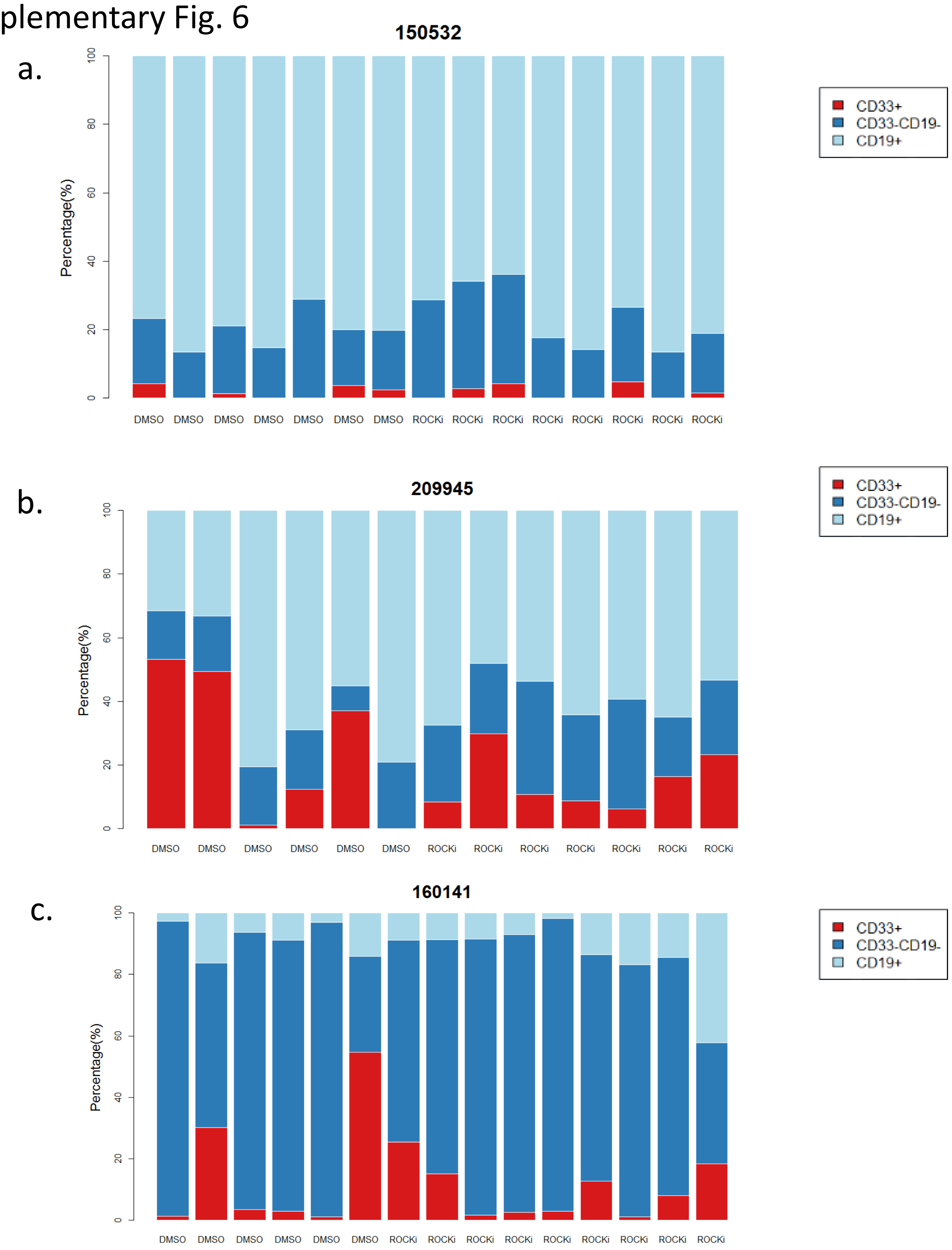

Supplementary Fig. 7

a.

**Hematoxicity of ROCKi in three CD34+ derived cell lineages**

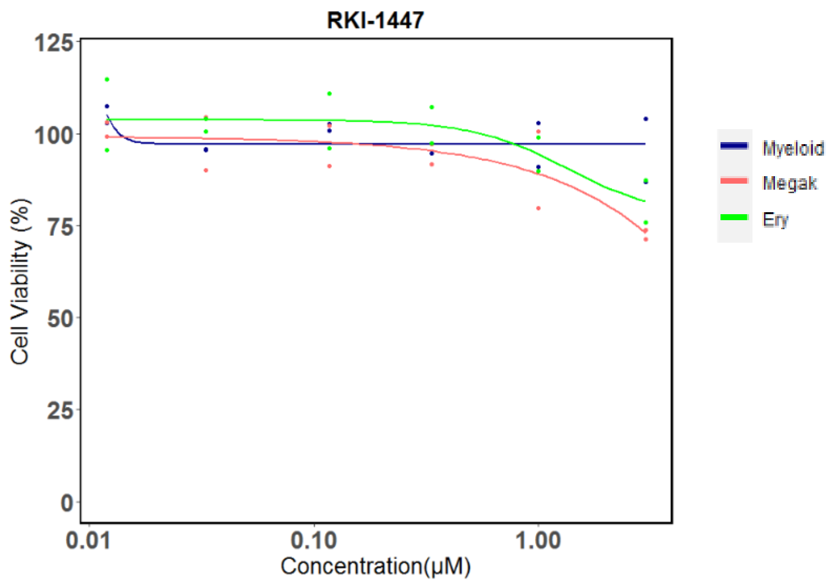


b.

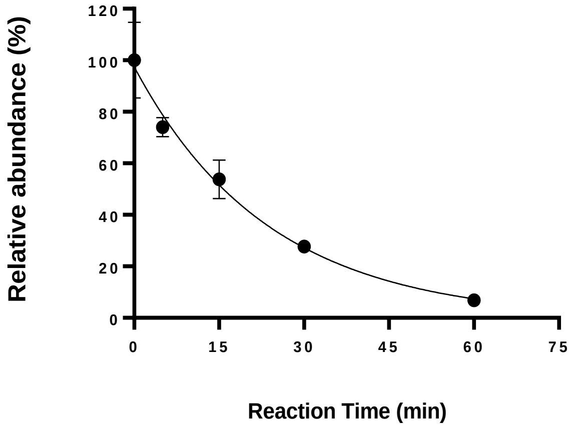


Half-life 16.45 min

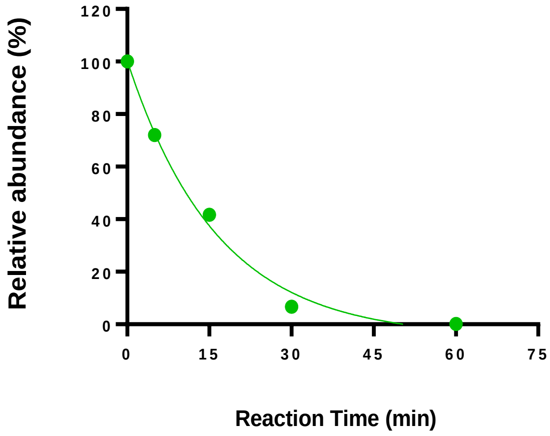


Half-life 11.42 min

Supplementary Fig. 8

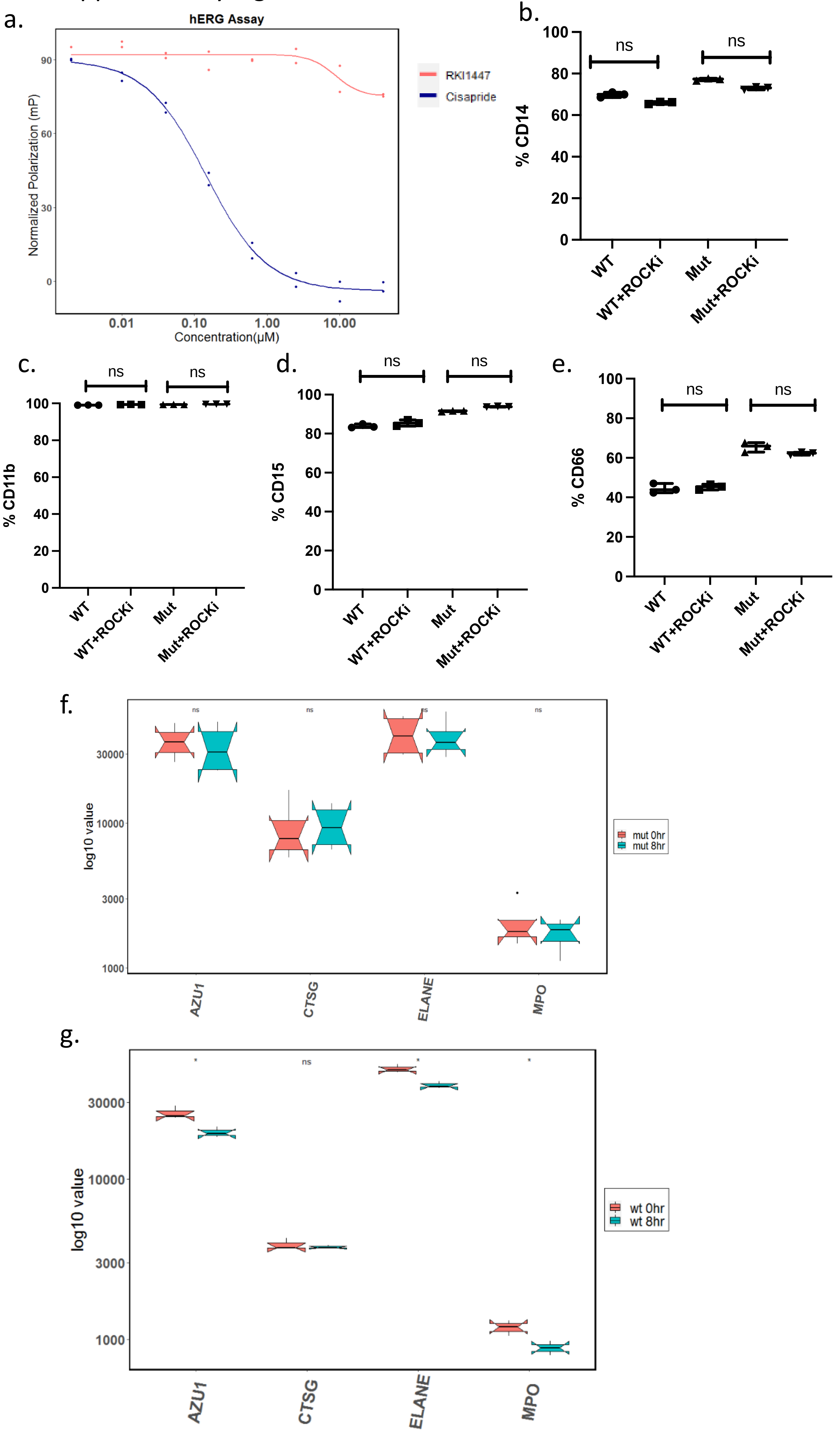

Supplementary Fig. 9

a.

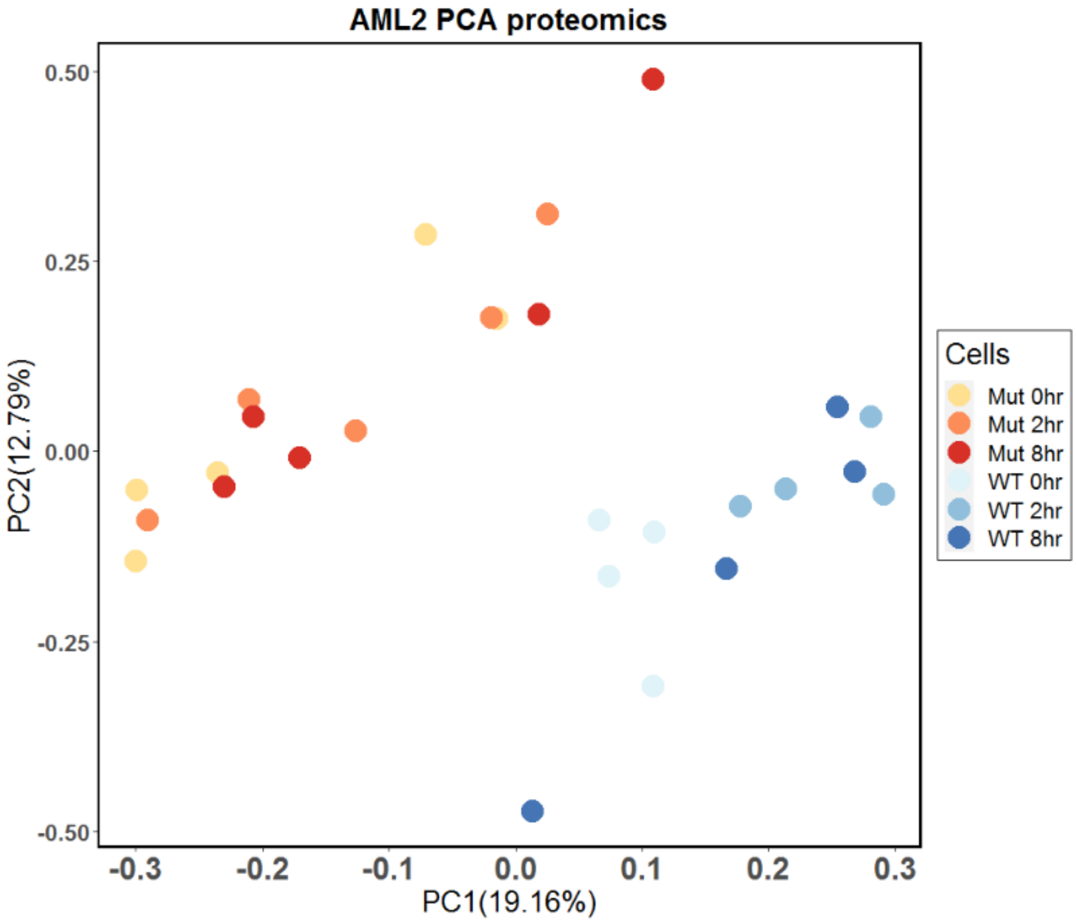


b.

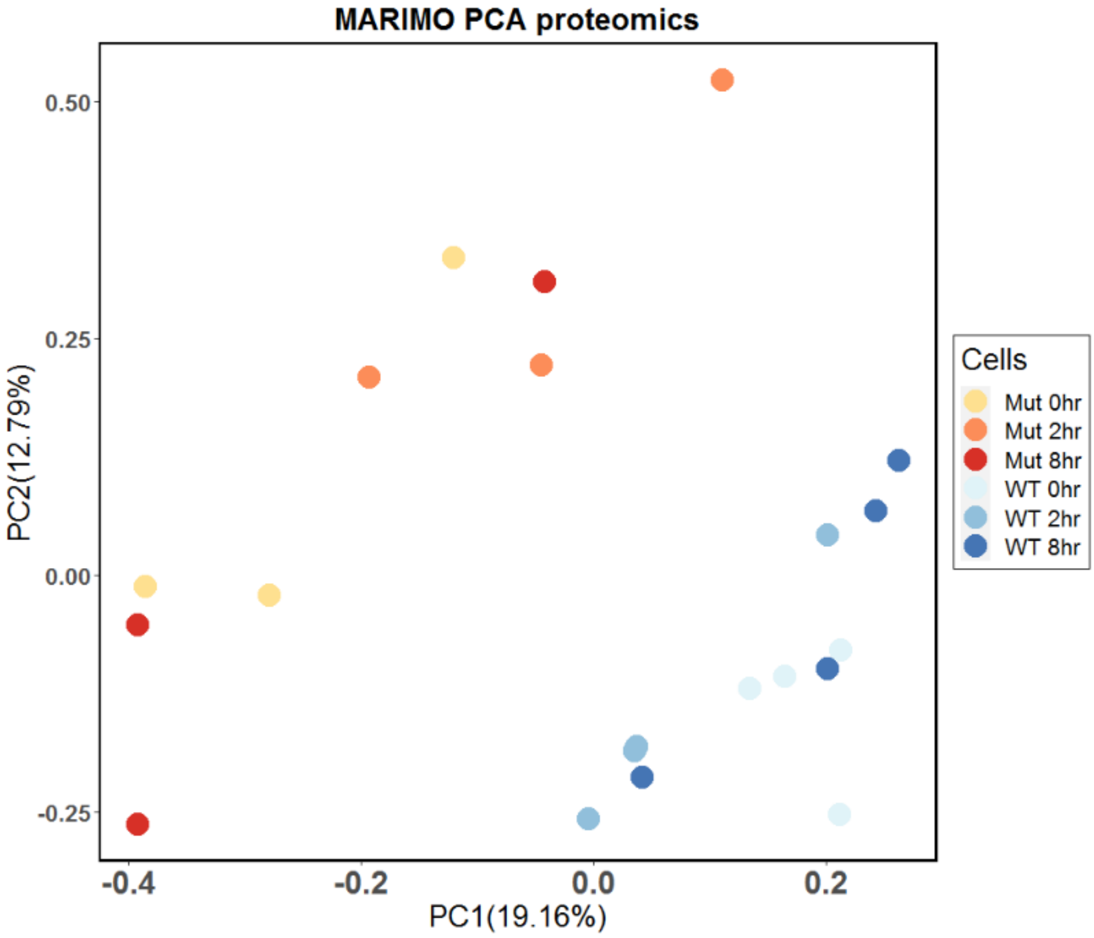

Supplementary Fig.10

#### MOLM14 WT

a.

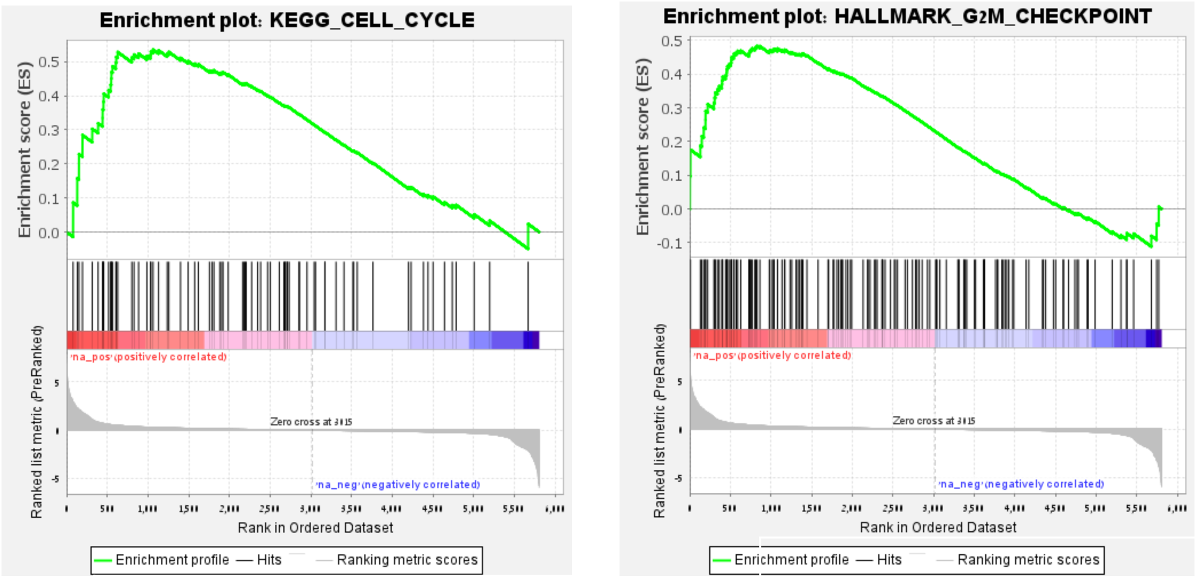


NES=1.4364926
Nominal p-value=0.02226345

NES=1.4271529
Nominal p-value= 0.010067114

#### MARIMO Mut

b.

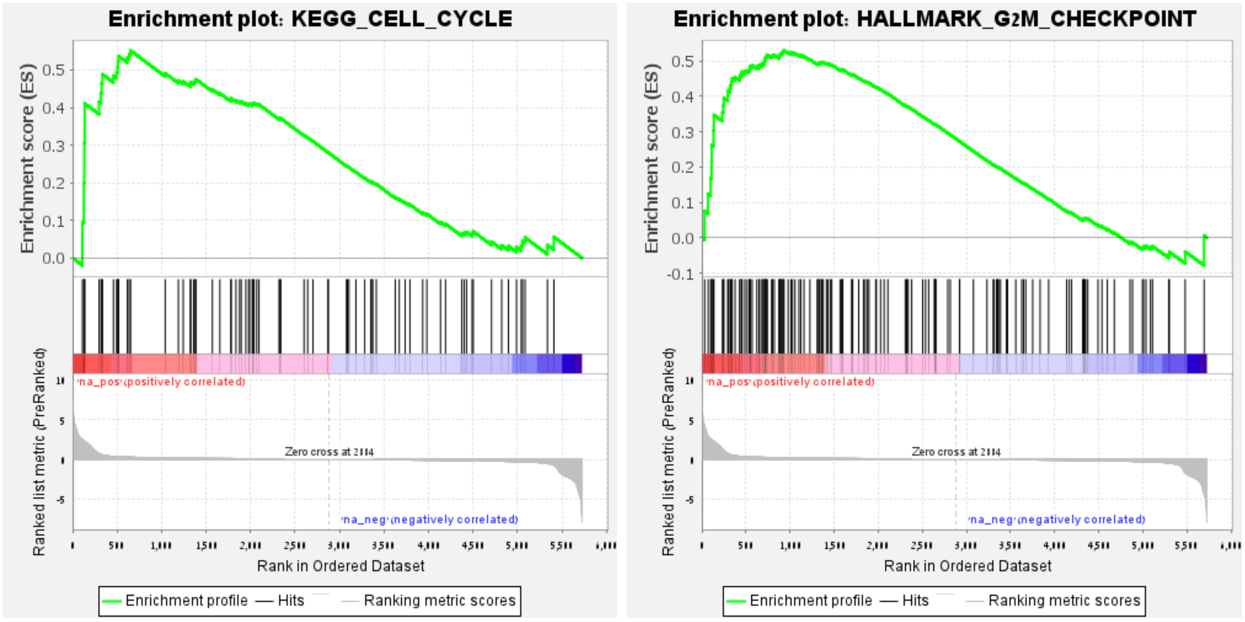


NES=1.3755981

Nominal p-value=0.046

NES=1.4247804

Nominal p-value=0.016

#### AML2 Mut

c.

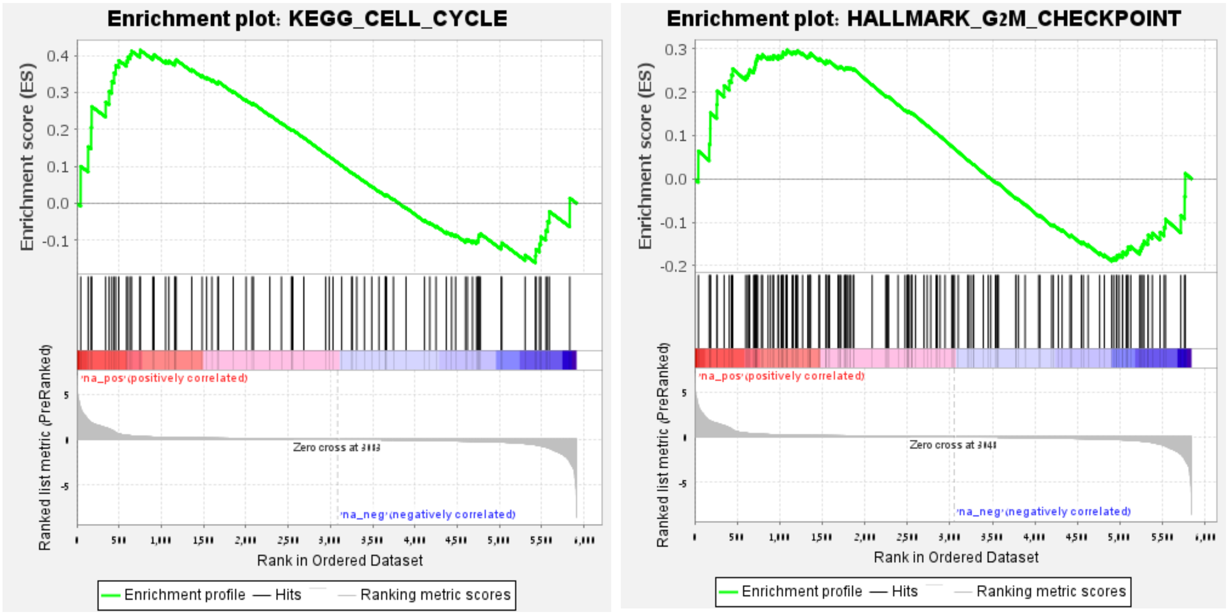


NES=1.0797888
Nominal p-value =0.32504147

NES=0.83830416
Nominal p-value=0.86292833

Supplementary Fig. 11

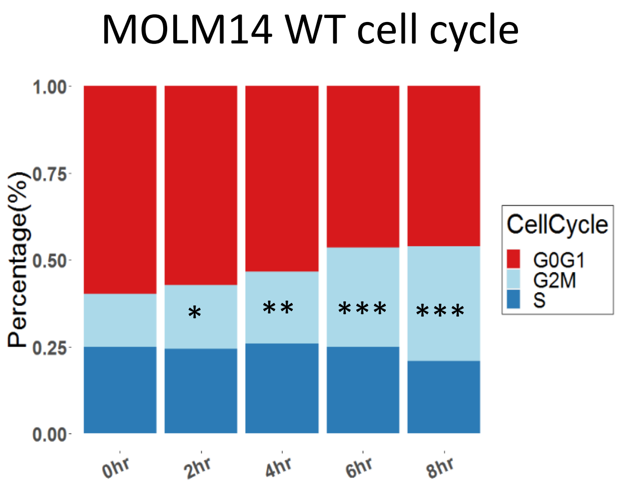

Supplementary Fig. 12

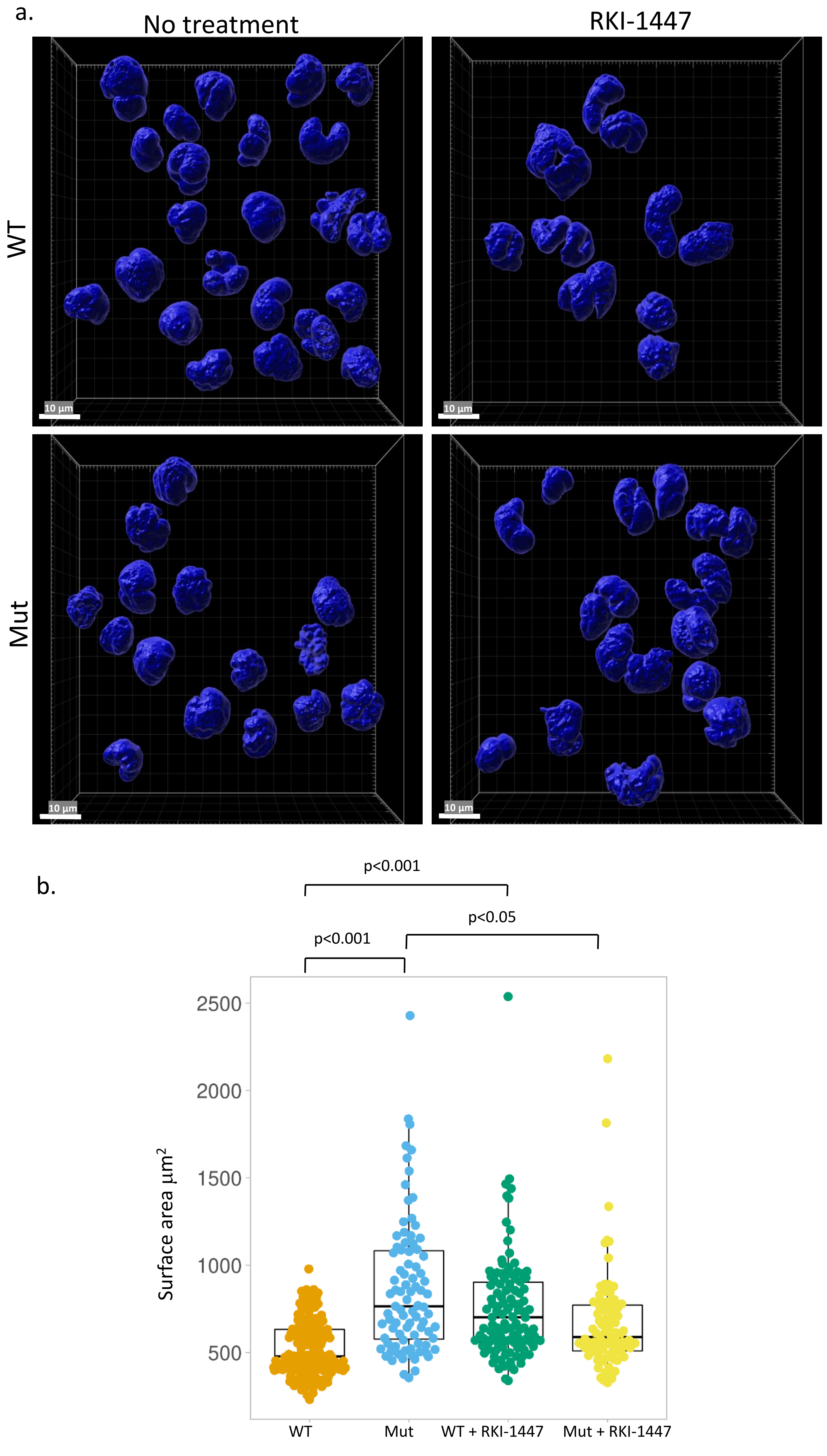

Supplementary Fig. 13

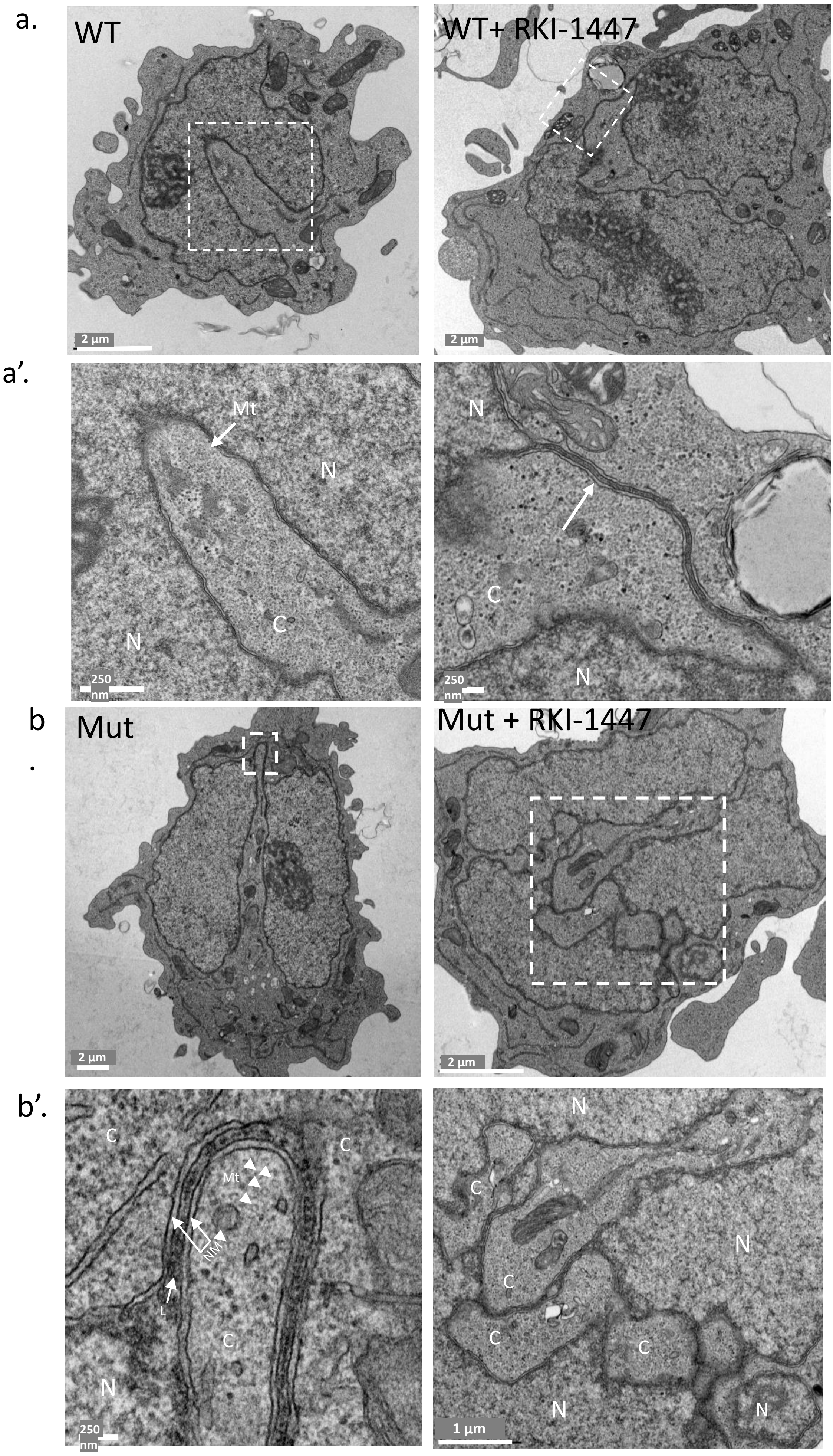

Supplementary Fig. 14

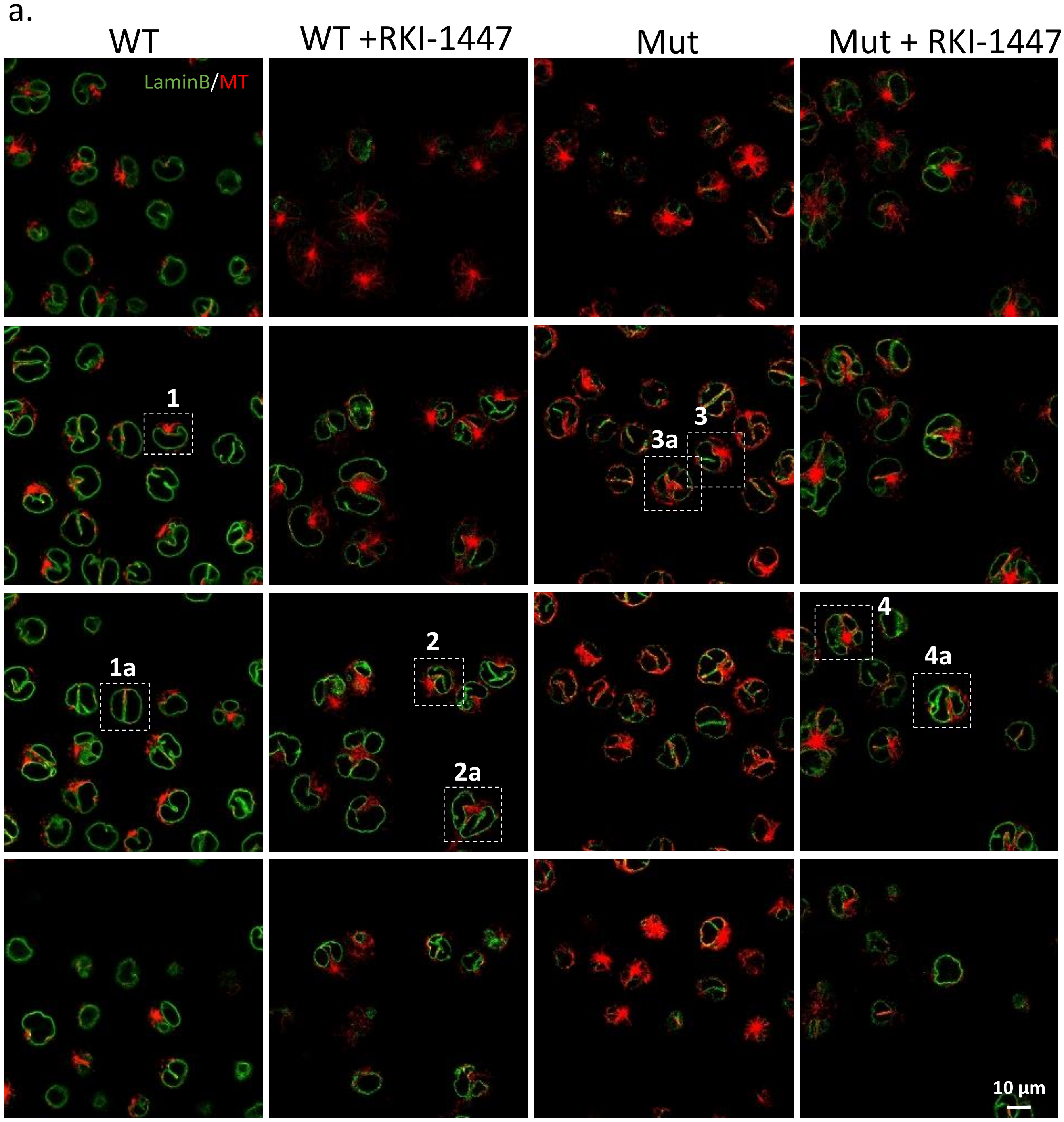


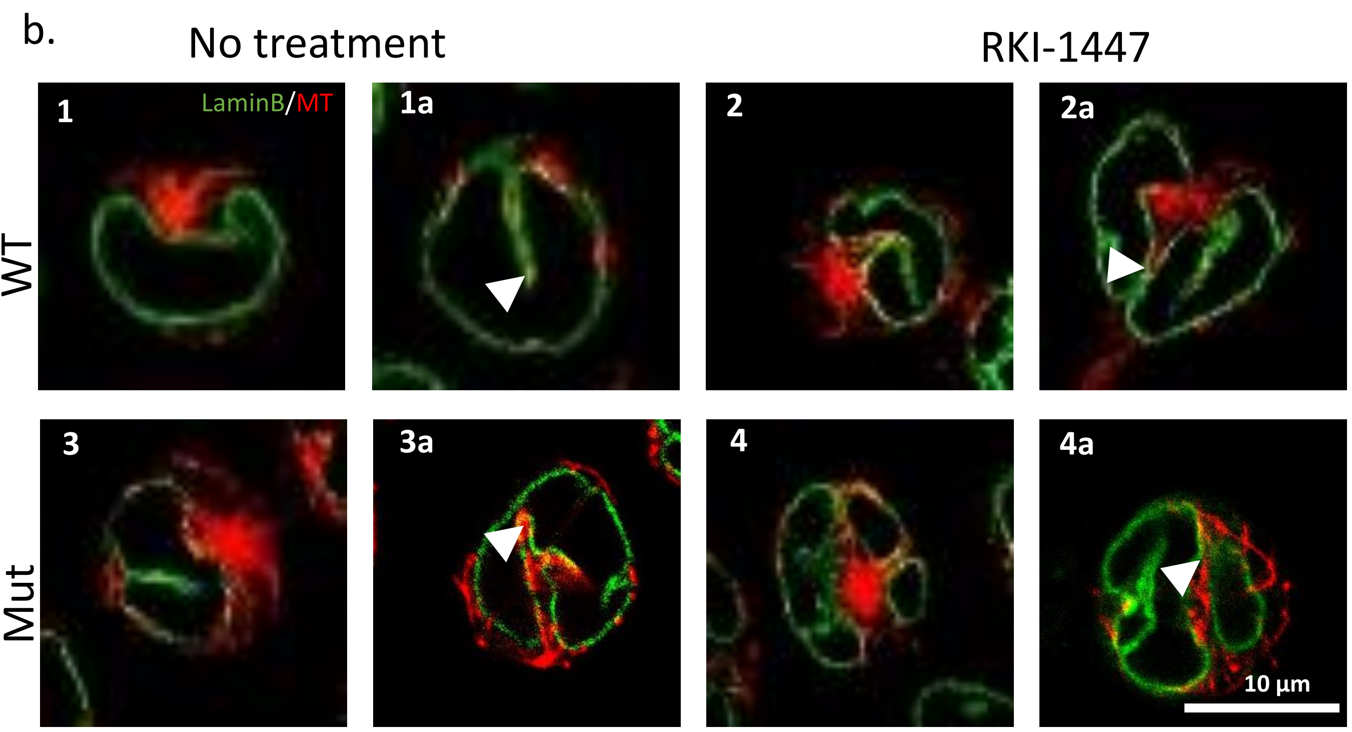

Supplementary Fig. 15

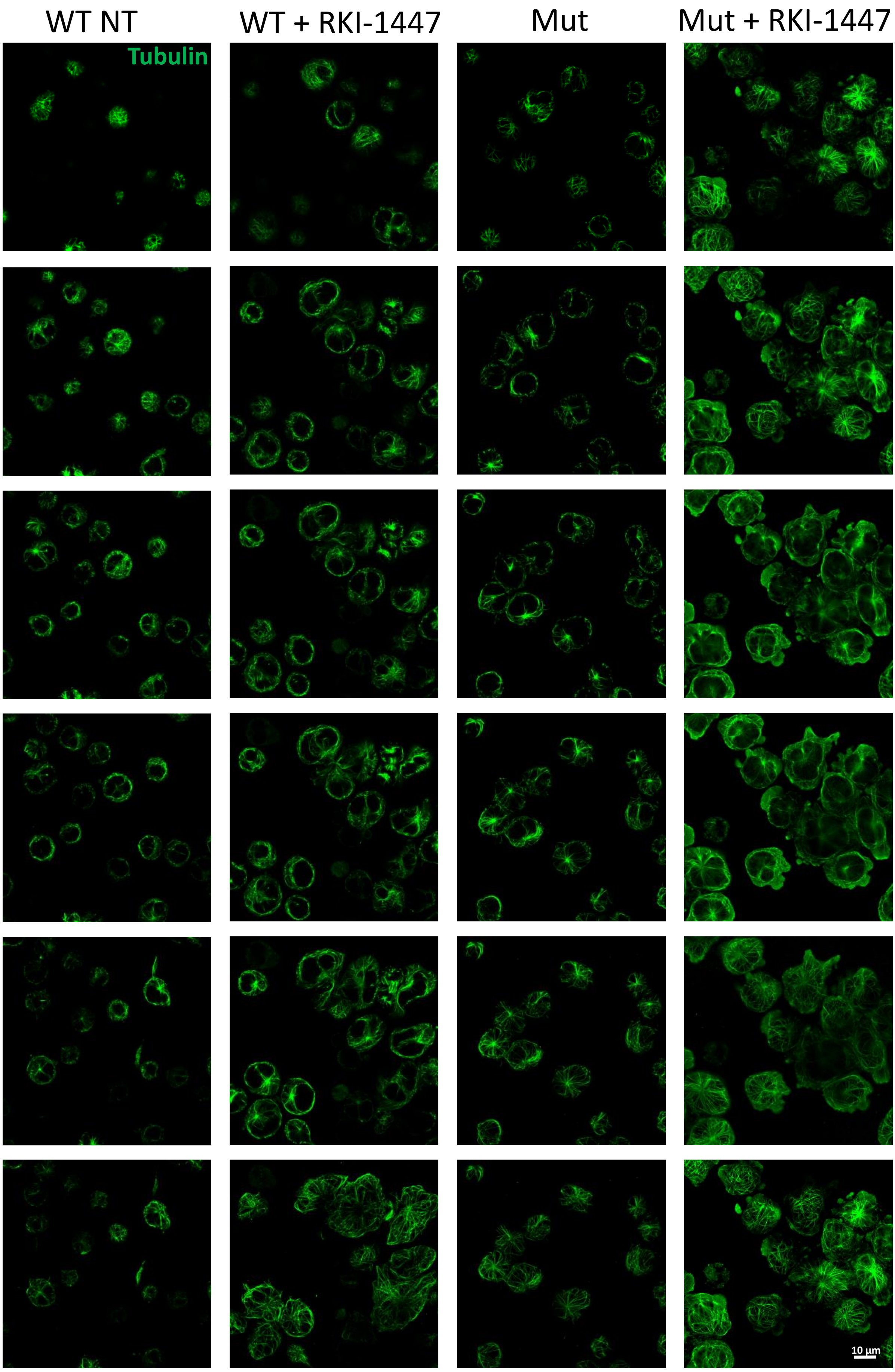

Supplementary Fig. 16

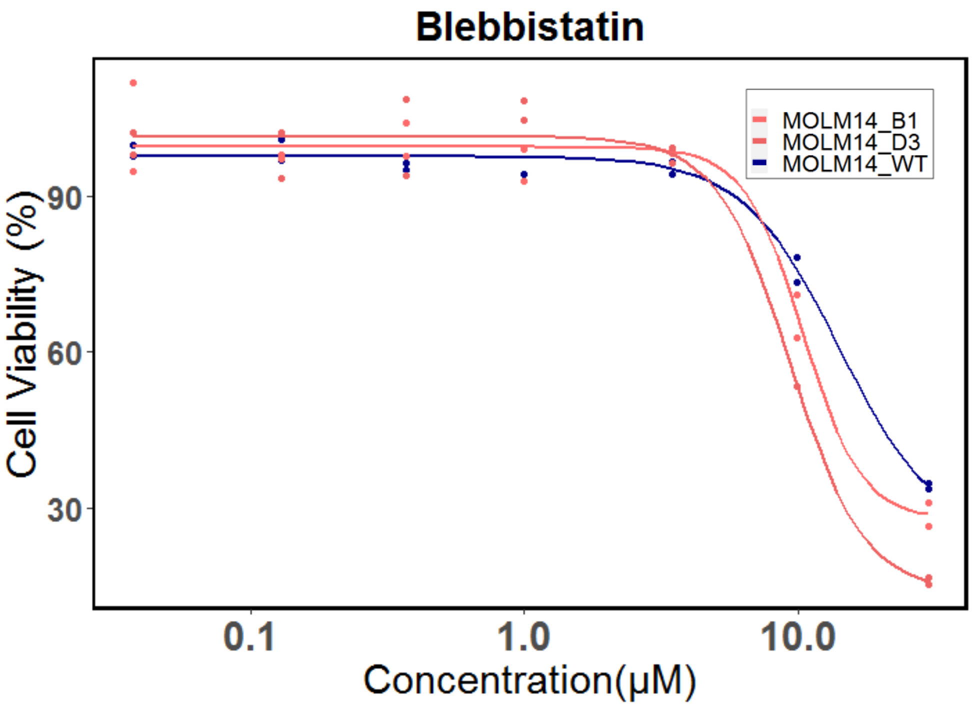

Supplementary Fig. 17

a.

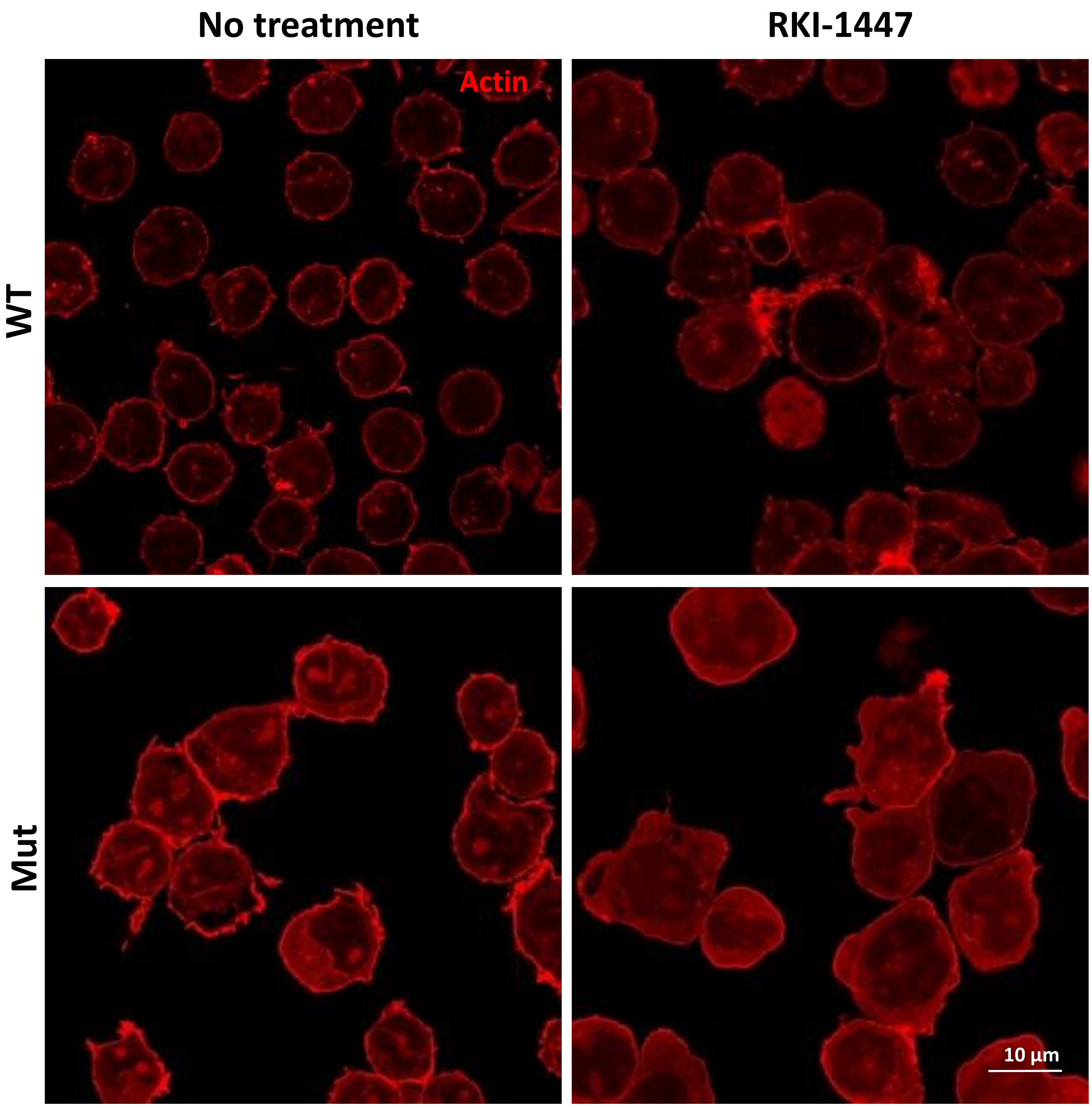


b.

| Sample | CY actin (MI) | NU actin (MI) | Ratio NU actin/CY actin |
|---|---|---|---|
| WT NT | 1809 | 933 | 0.51 |
| Mut NT | 3230 | 2788 | 0.86 |
| WT RKI-1447 | 1785 | 1208 | 0.68 |
| Mut RKI-1447 | 5773 | 3844 | 0.67 |
