## Supplementary information for "Rock inhibitors target *SRSF2* leukemia by disrupting cell mitosis and nuclear morphology"

**Supplemental Methods**

**Cell lines**

K562 cell line was purchased from the Stem Cell & Advanced Cell Technology Unit at the Weizmann Institute of Science and *SRSF2* Mut K562 cells was a generous gift from Omar Adel-Wahab’s lab; OCI-AML2, OCI-AML3, MOLM-14 and MARIMO cell lines were kindly donated by the Deininger/O'Hare Lab at the Huntsman Cancer Institute, University of Utah. All cells were cultured in RPMI-1640, 10% FBS and 1% P/S (01-100-1A, 04-007-1A and 03-031-1B, respectively, Biological Industries).

**Design of CRISPR guides and ssODN**

All oligo sequences were designed using Benchling Life Sciences R&D Cloud (https://benchling.com/). A 20bp sgRNA (GCGGCTGTGGTGTGAGTCCG) was designed for a SpCas9 (3’ side, PAM=NGG) system. The guide was designed to cut the SRSF2 exon 2, five bases downstream of the P95 SNP. To create the P95H mutation, a 120bp single-strand donor oligonucleotide (ssODN) was designed with as single mismatch so that it both altered the PAM motif as well as encoded for histidine instead of proline (GGGGCCGTGCTGGACGGCCGCGAGCTGCGGGTGCAAATGGCGCGCTACGGCCGCCACCCGGACTCACACCACAGCCGCCGGGGACCGCCACCCCGCAGGTACGGGGGCGGTGGCTACGGA).

**Cell transfection**

All cells were transfected in 20μl 16-well strips using a Lonza 4D-NucleofectorTM. Cells were sub-cultured 48hr prior to transfection at a concentration of 300,000 cells/ml. OCI-AML2 cells were transfected in SF solution (V4XC-2032, Lonza), 300,000 cells/rxn, using DN-100 program. MOLM-14 cells were transfected in SF solution, 1,000,000 cells/rxn, using DP-115 program. MARIMO cells were transfected in SF solution, 500,000 cells/rxn, using DN-100 program. K562 cells were transfected in SF solution, 200,000 cells/rxn, using FF-120 program. OCI-AML3 cells were transfected in SE solution (V4XC-1032, Lonza), 200,000 cells/rxn, using EO-100 program. The IDT Alt-R® CRISPR-Cas9 System Delivery of ribonucleoprotein complexes in HEK-293 protocol (v3.1) was used for reagent ratios. In brief, 2.1μl PBS (02-023-1A, Biological Industries), 1.2μl Alt-R® CRISPR-Cas9 sgRNA (100μM, IDT), 1.7μl Alt-R® S.p. Cas9 Nuclease V3 (61μM, IDT), for a total of 5μl/rxn. 1 μl Alt-R™ HDR Donor Oligo (200 μM, IDT) was added to each reaction. Following transfection cells were washed with medium, divided to two wells and cultured.

**DNA extraction**

Four days following transfection, bulk cells from one of the duplicate wells was centrifuged and DNA was extracted by way of lysis. 80µL of 50mM NaOH was added to each cell pellet, and heated at 99°C for 10 min. Cell lysate was then cooled on ice and 8µL of 1M Tris pH 8.0 was added.

**Next Generation Sequencing library preparation**

Libraries were prepared according to previously described methods^1^. In brief, primers for the amplification of SRSF2 P95 were designed, and 5’ adaptors were added to their sequence: (Fwd: CTACACGACGCTCTTCCGATCTctcagccccgtttacctg, Rev: CAGACGTGTGCTCTTCCGATCTctgaggacgctatggatg). Each PCR reaction contained 5µL of NEBNext® Ultra™ II Q5® Master Mix (M0544S, NEB), 0.5µL of each above primer (10µM), 4µL of cell lysate. The reaction was placed in a Eppendorf Mastercycler pro Thermal Cycler and the following protocol was initialized: 98°C for 30 sec; 33 cycles of 98°C for 10 sec and 65°C for 30 sec; 65°C for 5 min. The product of this reaction (‘PCR1’) was diluted 1:1000 and served as a template for the following reaction. Next, dual sequencing barcode were ordered according to the following formation: Fwd primer:

AATGATACGGCGACCACCGAGATCTACAC[Fw_Index_D5XX]ACACTCTTTCCCTACACGACGCTCTTCCG; Rev primer: CAAGCAGAAGACGGCATACGAGAT[Rev_Index_D7XX]GTGACTGGAGTTCAGACGTGTGCTCTTCCG. The second PCR reaction (‘PCR2’) contained 2.5µL of NEBNext® Ultra™ II Q5® Master Mix, 0.5µL nuclease-free water, 1µL of the diluted PCR1 template, and 1µL of the above barcode mix (2.5 µM). A total of 5µL were placed in the thermal cycler using the same protocol as above, for 28 cycles. The resulting PCR2 reaction was cleaned of any residual enzyme, nucleotides, and primer dimers traces, according to the recommended size selection protocol using AMPure XP SPRI magnetic beads (Beckman Coulter) at a volume ratio of x0.7.

**Single cell-sorting**

Four to seven days following the abovementioned transfection, bulk cells were stained with Propidium Iodide (556463, BD Pharmingen™) as per manufacturers’ instructions. Cells were sorted using a BD FACSAria™ III Cell Sorter, one cell per well, into Nunc™ Edge™ 96-Well Microplates (167425, Thermo Fisher). Cells were cultured for 2-4 weeks until colonies were visible, after which 100µL of cells and medium were transferred to Axygen® 96-well PCR plates (PCR-96-FS-C, Corning) and centrifuged at 400g for 5 minutes. The supernatant was decanted and DNA was extracted from cell pellets according to the lysis protocol described above. The resulting DNA was prepared according to the library preparation protocol described above, and proceeded to sequencing for genotyping of colonies.

**RNA-seq**

**RNA extraction**

RNA seq of all the cell lines are extracted with Qiuck-RNA MagBead (Zymo, R2132) according to the manual instruction.

**Library preparation**

Sequencing Libraries were prepared using INCPM mRNA Seq. NovaSeq 200 cycles reads were sequenced on 2 lane(s) of an Illumina novaseq. The output was ~54 million reads per sample.

**Gene expression analysis**

Gene expression pipeline was run with UTAP^2^ as following: Reads were trimmed using cutadapt (parameters: -a ADAPTER1 -a “A{10}” -a “T{10}” -A “A{10}” -A “T{10}” –times 2 -q 20 -m 25). Reads were mapped to genome hg19 using STAR v2.4.2a (parameters: –alignEndsType EndToEnd, –outFilterMismatchNoverLmax 0.05, –twopassMode Basic). The pipeline quantifies the Genecode annotated genes: hg19-genecode.genes.gtf. The annotation version and date are: description: evidence-based annotation of the human genome (GRCh37), version 19 (Ensembl 74). Counting was done using STAR. Further analysis is done for genes having minimum 5 read in at least one sample. Normalization of the counts was done using DESeq2 with the betaPrior set to True. Raw P values were adjusted for multiple testing using the procedure of Benjamini and Hochberg.

**Compound Libraries**

Three commercial libraries were used in the screening process: the Bioactive library (Selleck Collection, n=3727), the Kinom Set (n=187), and the Epigenetic chemical probe library (both from Structural Genomics Consortium, n=97).

**Cell viability assay**

Compounds were dispensed in 384-well plates using an ECHO® 555 liquid handler (Labcyte) and sealed. On day of experiment, the concentrations of isogenic and respective wildtype cell lines were counted using a Countess™ II FL (InvtrogenTM) and re-suspended at 40,000 cell/ml. 50µL of medium was dispensed using a Multidrop™ Combi Reagent Dispenser (Thermo Fisher), bringing the total number of cells in each well to 2000. Cells were incubated for 48 hours. On day of measurement, plates were centrifuged and supernatant was removed using a Washer/Dispenser II (GNF Systems). A Washer/Dispenser II was then used to dispense CellTiter-Glo® (G7572, Promega) as per the manufacturer’s instructions. The luminescence signal was then measured using a PHERAstar® FSX (BMG Labtech) and results were analyzed using Genedata Screener®. The viability of treated cells was normalized to a vehicle control, contained on each plate.

**siRNA**

**Preparation of siRNA stock**

ON-TARGETplus human ROCK1 siRNA (L-003536-00-0005 NM_005406) and ON-TARGETplus Human ROCK2 siRNA (L-004610-00-0005 NM_001321643 NM_004850) and ON-TARGEplus non-targetin Pool were purchased from Dharmacon. SiRNA(5nmol) were resuspended in 50 µl RNase-free water or siRNA suspension buffer to obtain a 100µM siRNA stock solution. Heat the tube to 90°C for 1 min Incubate at 37°C for 60 min and stock at -20°C.

**Transfection**

Cells were sub-cultured 48hr prior to transfection at a concentration of 300,000 cells/ml. Cells were transfected in 20μl 16-well strips using a Lonza 4D-NucleofectorTM. Cells were transfected in SF solution, 500,000 cells/rxn, using DP-115 program. 250nM siRNA pool targeted *ROCK1* / *ROCK2* and non-targeted siRNA pool added to different reaction. Following transfection cells were divided to two wells and cultured. 48 hours after transfection, the number of the viable cells are counted manually and normalized to non-targeted well.

**In vivo**

NSG mice (n=5-10/sample) were injected with half million to five million CD3 depleted AML cells from the peripheral blood of six patients with *SRSF2* mutated AML intra-femorally. On day 35 the animals were randomized to RKI-1447 or a carrier control. RKI-1447 is given i.p. 50mg/kg. The weight of NSG mice was measured before and after drug administration. On day 56 mice were sacrificed and analyzed for human engraftment by flow cytometry and the presence of *SRSF2* Mutations of the CD45+ human engrafted cells are sequenced with target sequence of P95 region. Library preparation and sequencing are performed as we mentioned at the previous section.

**Flow cytometry**

Primary human samples and cells extracted from mice bone marrows were stained with the antibodies shown in Table 1.

**Table 1**: Antibodies and viability staining used for flow cytometry

| Antibody Manufacturer Clone/Catalogue number Titer | | | |
| --- | --- | --- | --- |
| CD45-BV510 | BioLegend, San Diego, CA, USA | HI30 | 1:200 |
| CD33-APC | BD Biosciences, San Jose, CA, USA | WM53 | 1:100 |
| CD34-APC Cy7 | BioLegend, San Diego, CA, USA | 581 | 1:100 |
| CD15-BV421 | BioLegend, San Diego, CA, USA | W6D3 | 1:100 |
| CD38-PE Cy7 | BioLegend, San Diego, CA, USA | HIT2 | 1:100 |
| CD3-FITC | BD Biosciences, San Jose, CA, USA | UCHT1 | 1:100 |
| CD19-PE | BD Biosciences, San Jose, CA, USA | HIB19 | 1:200 |
| PI | BD Biosciences, San Jose, CA, USA | 556463 | 1:100 |

**DNA extraction of the grafts**

The human CD45+ grafts of NSG mice are separated with EasySep™ Human CD45 Depletion Kit II (Catalog 18259) for each mouse and then DNA of the cells are extracted immediately with QIAamp® DNA Micro kit (Qiagen 157032361) according to the manual instruction.

**Toxicity**

**Hepatoxicity**

**Cell lines**

THLE-2 (CRL-2706™) were purchased from ATCC and were grown in the BEGM Bullet Kit (CC-3170) from Lonza. Besides the additives contained in the kit, the medium was further supplemented with 5 ng/mL EGF (Sigma), 70 ng/mL phosphoethanolamine (Sigma) and 10% FBS. The plates for the THLE-2 needed to be pre-coated with a mixture of 0.01mg/mL fibronectin, 0.05mg/mL of PureCol™ EZ Gel Solution (Sigma) and 0.01mg/mL of BSA dissolved in BEBM medium (Lonza). The coating medium was aspirated before seeding.

**Assessment of compound toxicity in THLE-2**

THLE-2 cells were exposed to the compounds in a 9-point, 2-fold dilution dose response series with a 100uM as upper limit, for 72 hours. Following the 72 hours of exposure to the test compounds, cell viability was determined by measuring the concentration of cellular ATP (CellTiter Glo, Promega). The luminescence signal was measured on a Pherastar FS multi-mode plate reader (BMG Labtech). Each data point was run in triplicate.

**The Predictor™ hERG Assay**

The assay was performed in accordance to the manufacture protocol (Invitrogen, #PV5365). Shortly, a small volume medium binding 384-well plates (Greiner, #784076), were preplated with RKI1447, Cisapride (positive control), and DMSO (vehicle) using an ECHO® 555 liquid handler (Labcyte). Additional controls – E-4031 (kit positive control), blank, free tracer and negative control were added in accordance to the manufacture protocol. The FP signal was measured using 540/590 fluorescent polarization filter of PHERAstar® FSX (BMG Labtech) and results were analyzed using Genedata Screener®. The polarization (mP) signal was normalized to negative/positive controls.

**Metabolic stability assay**

2 mM Compounds in DMSO are prepared and stored at -20 degrees. Pooled Human Liver Microsomes (Mixed Gender Sigma M0317) are diluted to 0.6 mg/ml in 100 mM potassium phosphate buffer (pH 7.4) with 1.0 mM NADPH (Sigma N5130).  Compounds (1 µM final) are incubated with microsomes and NADPH at 37 degrees for a time course using thin-walled PCR tubes and a thermocycler set to constant temperature. At each time-point, 5 µl aliquots are removed to 25 µl ice cold Acetonitrile containing 10 µg/ml Kaempherol (Sigma K0133) as an internal control. Centrifuge 2500xg for 15 minutes, add 20 µl ice-cold DDW and repeat centrifuge. Transfer to 384-well plate (Waters 186002631), seal with aluminum seal and store at -80 degrees. Compounds are quantified by LC/MS. Data is normalized to percent of input compound in buffer without microsomes and time-course is plotted with Graphpad Prism.

**Hematotoxicity**

**CD34+ cells separation**

Enrichment of human CD34+ cells from PBMC of healthy individual was performed according to the manufacturer’s instructions (Miltenyi Biotech, Bergisch Gladbach, Germany). Seed CD34+ cells in 384-well plate at a density of 1000 cells per well in 70ul SFEMII medium with different cytokine cocktail (Table2) for myeloid, erythroid and megakaryocytic lineages, culture in an incubator for 7 days at 37 degrees in a humidified atmosphere containing 5% CO2.

**Table2. Cytokine cocktails for myeloid, erythroid and megakaryocytic lineages**

| **Cytokines** | **Manufacturer** | **Cat** | **myeloid** | **erythroid** | **megakaryocytic** |
| --- | --- | --- | --- | --- | --- |
| rh Flt-3 Ligand | GenScript | Z02926-50 | 20(ng/ml) |  |  |
| rh Stem Cell Factor | GenScript | Z02692-50 | 100(ng/ml) | 100(ng/ml) | 100(ng/ml) |
| rh IL-3 | GenScript | Z03156-50 | 10(ng/ml) | 10(ng/ml) |  |
| rh IL-6 | GenScript | Z03034-50 | 50(ng/ml) |  | 50(ng/ml) |
| rh GM-CSF | GenScript | Z02983-50 | 20(ng/ml) |  |  |
| rh Erythropoietin | GenScript | Z02975-50 |  | 3(ng/ml) |  |
| rh IL-9 | GenScript | Z03013-50 |  |  | 10(ng/ml) |
| rh Thrombopoietin | peprotech | 300-18-10ug |  |  | 100(ng/ml) |

On Day 7, cells were treated with different concentrations of RKI-1447. On Day 9, the viability of cells is detected with CTG assay and the duplicate plate was transferred to 96-well plate and stained with the antibodies (**Table 3**) for flow cytometry to determine the fraction of differentiated cells after treated with RKI-1447.

**Table3.** Antibodies used for flow cytometry of hematotoxicity

| **Lineages/cells** | **Antibodies** | **Manufacturer** | **Cat** | **dilution** |
| --- | --- | --- | --- | --- |
| Megakaryocytic | CD42b-APC | BioLegend | 303912 | 1:100 |
| Myeloid | CD11b/MAC-1-APC-Cy7 | BD | PMG-557754 | 1:100 |
|  | CD66b- BV421 | BD | 562940 | 1:100 |
|  | CD15-BV650 | BD clone | 564232 | 1:100 |
|  | CD14-PE-Cy7 | BD clone | 557742 | 1:100 |
| Erythroid | CD71-FITC | BD | 347513 | 1:100 |
|  | CD235a-PE | BioLegend | 349105 | 1:100 |
| Stem/Progenitors | CD34-pecy5.5 | eBioscience | 46-0349-42 | 1:100 |

**iPSCs**

SRSF2 P95L and isogenic WT iPSCs were generated and differentiated into HSPCs as previously described^3,4^. iPSC-derived HSPCs were plated at a density of 8k cells per well in 96-well plates and treated with RKI-1447 (3μM) or DMSO for 48 hours. Viability was measured with CellTiter-Glo® (G7572, Promega) according to the manufacturer’s instructions.

**Mass spectrometry**

**Sample preparation, LC and MS**

The samples were subjected to tryptic digestion using an S-trap. The resulting peptides were analyzed using nanoflow liquid chromatography (nanoAcquity) coupled to high resolution, high mass accuracy mass spectrometry (Fusion Lumos). Each sample was analyzed on the instrument separately in a random order in discovery mode.

**Data processing**

Raw data was processed with MaxQuant v1.6.6.0. The data was searched with the Andromeda search engine against the human proteome database appended with common lab protein contaminants and the following modifications: Carbamidomethylation of C as a fixed modification and oxidation of M and protein N-terminal acetylation as variable ones. The LFQ (Label-Free Quantification) intensities were calculated and used for further calculations using Perseus v1.6.2.3. Decoy hits were filtered out, as well as proteins that were identified on the basis of a modified peptide only. The common contaminates are labeled with a ‘+’ sign in the relevant column. The LFQ intensities were log transformed and only proteins that had at least 2 valid values in at least one experimental group were kept. The remaining missing values were imputed.

**PCA**

Proteins labeled with a “+” sign are removed and then proceeded with principal component analysis in R with prcomp function.

**GSEA analysis**

The gene names were ranked based on fold change of the proteomics between different conditions and analyzed using the GSEA software version 4.1.0. Number of permutations: 1000. Gene sets database h.hall.v7.4.symbols.gmt [Hallmarks] and c2.cp.kegg.v7.4.symbols.gmt [Curated] were used.

**Cell Cycle**

Cells were collected and washed twice with PBS. Cells were suspended with 0.5 ml PBS and 4.5 ml of ice-cold EtOH 70% were added. Cells were mix well in the tube and kept on ice for at least 2 hours.

Centrifuge the ethanol-suspended cells. Decant ethanol thoroughly. Resuspend the cell pellet in 5 ml PBS, and centrifuge. Resuspend cell pellet in 1 ml PI/Triton X-100 staining solution with RNase. Keep it either 15 min at 37°C or 30 min at room temperature.

Set up and adjust the flow cytometer for excitation with blue light and detection of PI emission at red wavelengths. Measure cell fluorescence in the flow cytometer. Use the pulse width–pulse area signal to discriminate between G2 cells and cell doublets and gate out the latter. Read PI at linear scale. Analyze the data using DNA content frequency histogram deconvolution software.

**Propidium iodide (PI)/Triton X-100 staining solution with RNase A**

To 10 ml of 0.1% (v/v) Triton X-100 (Sigma 93443-100ML) in PBS add 2 mg DNase-free RNaseA (Sigma R6513-50MG) and 200 μl of 1 mg/ml PI (Sigma P4170-25MG). Prepare freshly.

**Immunofluorescent cell labeling**

Suspension of MOLM14 cells, both wild type (WT) or caring the B1 mutation in SRSF2, at the density of 300,000 cells/ml were cultured for 24 hours either in fresh RPMI 10%FBS 1%P/S medium, or in medium containing 0.5 µM RKI-1447. The cells were then plated onto optical plastic bottom 24 well dishes (IBIDI, Gewerbehof Gräfelfing). The cells were allowed to adhere to the well for 10 minutes^5^ and then washed with PBS and fixed-permeabilized for 1 minute with cytoskeleton stabilizing buffer:10 mM MES, 150 mM NaCl, 5 mM EGTA, 5 mM MgCl2, and 5 mM glucose, pH 6.1 containing 0.5% glutaraldehyde (Electron Microscopy Science) and 0.25% TrytonX100, and postfixed with 1% glutaraldehyde for 10 minutes ^6^ .The cells were then washed three times with PBS and treated for 10 minutes with 1mM sodium borohydride to reduce non-specific cellular autofluorescence.

Cells were labeled with either mouse monoclonal antibodies against α-tubulin (clone DM1A dilution 1:500, Sigma) or with rabbit monoclonal antibodies against Lamin B1 (clone 6K22, dilution 1:100, Sigma). Secondary antibodies used in these experiments were Alexa-488 conjugated goat-anti mouse or anti-rabbit antibodies 1:200 dilution (Invitrogen) was used. Actin was labeled using TRITC- Phalloidin (Sigma) 5ug /ml; and nuclei were labeled with DAPI (Sigma).

**Confocal microscopy and image analysis**

The cells were imaged on an inverted Olympus confocal FluoView FV1000 IX81 confocal laser-scanning microscope, equipped with a Diode laser with 405 nm emission, an Argon-ion laser with 488 nm emission and a Helium-Neon laser with 560 nm emission beam for dye excitation. Confocal microscopy scanning was sequential with a 4 μs/pixel dwell time. Emission signals were collected using PMTs. Images were sampled at 1600 by 1600 pixels with a bit depth of 12, using a 60x oil immersion objective (NA 1.35), and a 2X software zoom. The pinhole size used was 2 AU (119 µm). Z stacks were acquired with a step size of 0.3 μm. The images were visualized by Fluoview Olympus confocal software and afterwards processed by Imaris or ImageJ software.

A second set of confocal images were acquired, using an inverted Leica SP8 STED3X confocal microscope, equipped with internal Hybrid (HyD) detectors and Acusto-Optical Tunable Filter (Leica microsystems CMS GmbH, Germany) and The White light laser (WLL) excitation laser ranging from 470 - 670 nm. Dye excitation was performed using the white and diode laser. Emission signal was collected using an internal HyD detector. Imaging was acquired with STED-objective HCX PL APO 93x/1.30 GLYC STED White motCORR. The scanning was sequential and bi-directional with a scanning speed of 700 Hz. Images were sampled at 2328 by 2328 pixels with a bit depth of 16 and a pixel size of 0.042/0.042/0.35 µm for x/y/z dimensions. The zoom was set to 0.75, the line average 2 with a pinhole size of 2 AU (119 µm). Z stacks were acquired using the galvo stage, with a step size of 0.38 µm. The deconvolution of the Leica software was applied and the acquired images were visualized using LASX software (Leica Application Suite XLeica microsystems CMS GmbH).

Three-dimensional reconstruction and image analysis were carried out using the Imaris software, Oxford instruments (Bitplane) package, version 9.7.1. For the 3D nuclear shape analysis, their surfaces were rendered by manual thresholding that was kept constant, and the data such as area and volume was exported to excel files for statistical analysis.

The actin intensity was analyzed using ImageJ (Fiji) by masking with an Otsu threshold, for the actin (based on the TRITC phalloidin staining) and the nuclei (using DAPI staining). The actin mask was eroded to exclude the cortical actin. Thereafter the masks were subtracted to create a cytosolic mask. The intensity was measure by selection of the different masks; nuclei and cytosolic and the results were summed in the table (**Supplementary Fig.17b**). The images for the analysis are 16-bit images from the Leica confocal SP8 microscope.

**Transmission electron microscopy**

Cells were fixed for 1 hour with 4% paraformaldehyde, 2% glutaraldehyde in 0.1 M cacodylate buffer containing 5 mM CaCl2 (pH 7.4), postfixed in 1% osmium tetroxide supplemented with 0.5% potassium hexacyanoferrate trihydrate and potassium dichromate, stained with 2% uranyl acetate in double distilled water for 1 hour, dehydrated in graded ethanol solutions and embedded in epoxy resin. Ultrathin sections (70 nm) were obtained with a Leica EMUC7 ultramicrotome and transferred to 200 mesh copper transmission electron microscopy grids (SPI). Grids were stained with lead citrate and examined with a Tecnai SPIRIT transmission electron microscope (Thermo Fisher Scientific). Digital electron micrographs were acquired with a bottom-mounted Gatan OneView camera.

**Cytoskeleton drugs**

34 cytoskeleton drugs including myosin II, microtubule and actin modulating compounds were used to perform dose-response assay on MOLM14 WT and *SRSF2* Mut cells at concentration of 0.037μM, 0.129μM, 0.369μM, 0.997μM and 29.900μM in duplicates. Cell viability were detected as described above.

**Splicing analysis**

We used rMATS v4.0.2 to assess the differential splicing landscape embedded in RNA-seq data. All of the aberrant splicing events used the cut off FDR<0.05 and ΔPSI >0.1 (10%).

**Statistical analysis**

In all mice engraftment figures comparison between medians was performed using the Mann−Whitney U test with FDR correction for multiple hypothesis testing.

The p-value of the overlap of aberrant exon usage gene of all our SRSF2 Mut cell lines and *SRSF2* mut BeatAML database were analyzed with the following code in R:

hygeo6<-function(N,m,a,b,c,g,h,w){

#N=11027# total number of genes for selection

#m=14 # number of overlaps in all three sets

#a=1332# number of aberrant spliced genes in OCI-AML2 *SRSF2* Mut

#b=1295# number of aberrant spliced genes in MARIMO *SRSF2* Mut

#c=1025# number of aberrant spliced genes in MOLM14 *SRSF2* Mut

#g=1579# number of aberrant spliced genes in OCI-AML3 *SRSF2* Mut

#h=1723# number of aberrant spliced genes in K562 *SRSF2* Mut

#w=488# number of aberrant spliced genes in BeatAML *SRSF2* Mut

d<-data.frame(0:min(a,b,c,g,h,w),rep(0,min(a,b,c,g,h,w)+1))

for (i in 1:10000){

A=sample(1:N,size=a,replace=FALSE)

B=sample(1:N,size=b,replace=FALSE)

C=sample(1:N,size=c,replace=FALSE)

G=sample(1:N,size=g,replace=FALSE)

H=sample(1:N,size=h,replace=FALSE)

W=sample(1:N,size=w,replace=FALSE)

e<-table((C %in% A)&(C %in% B)&(C %in% G)&(C %in% H)&(C %in% W))["TRUE"]

if(!complete.cases(e)){

e<-0

}

d[e+1,2]<-d[e+1,2]+1

}

colnames(d)<-c("Intersect","p-value")

d[,2]<-d[,2]/10000

p<-sum(d[(m+1):(min(a,b,c,g,h,w)+1),2])

print(d)

return(p)

}

hygeo6(11027,14,1332,1295,1025,1579,1723,488)

**Supplementary Tables**

Supplementary Table 1 - Targeted amplicon sequencing of all *SRSF2* Mut cell lines.

Supplementary Table 2 - Aberrant splicing events of *SRSF2* Mut cells lines

Supplementary Table 3 - List of exon skipped (SE) genes of *SRSF2* Mut cell lines and BeatAML database

Supplementary Table 4 - HTDS of 3988 chemical compounds in a single dose (10μM)

Supplementary Table 5 - Dose response analysis and IC-50 of 44 compounds

Supplementary Table 6 - Dose response analysis and IC-50 of four ROCK inhibitors on MOLM14, AML2 and MARIMO cell lines

Supplementary Table 7 - Clinical and molecular data of *SRSF2* mut samples and CD34+ sample

Supplementary Table 8 - Flow cytometry of *SRSF2* Mut and healthy CD34+ xenograft model

Supplementary Table 9 - Liver toxicity of RKI-1447

Supplementary Table 10 - Quantification of 3D rendering of nucleus of *SRSF2* WT and Mut MOLM14 cell lines before and after exposure to RKI-1447

**Supplementary Fig. 1: The cell proliferation growth curves of WT and *SRSF2* Mut cell lines**. **a**, OCI-AML2. **b**, OCI-AML3. **c**, MARIMIO. **d**, K562. Technical duplicates of each line and each day.

**Supplementary Fig. 2: Dose-response curve of WT and *SRSF2* Mut OCI-AML2 and MARIMO cell lines of four different ROCKi compounds.** **a-d**, dose-respond curve of *SRSF2* WT and *SRSF2* Mut OCI-AML2 cells (AML2_E9, AML2_B7, AML2_F6, AML2_B3, AML2_H1) on (**a**)RKI-1447, (**b**) GSK429286A, (**c**) GSK180736A and (**d**) Y-39983 at concentration of 0.037μM, 0.129 μM, 0.369μM, 0.997μM, 29.900μM. **e-h**, dose-respond curve of *SRSF2* WT and *SRSF2* Mut MARIMO cells (MARIMO_A10, MARIMO_A12, MARIMO_A2, MARIMO_C1, MARIMO_D12, MARIMO_E3, MARIMO_G4) on (**e**)RKI-1447, (**f**) GSK429286A, (**g**) GSK180736A and (**h**) Y-39983 at concentration of 0.037μM, 0.129μM, 0.369μM, 0.997μM, 29.900 μM.

**Supplementary Fig. 3: Engraftment of MOLM14 *SRSF2* Mut cells in NSG mice.** NSG mice were treated with RKI-1447 (50mg/kg) (N=4) or vehicle (N=3) for 3 weeks after 5-week transplantation of one million *SRSF2* Mut molm14 cells (i.v.). Mice were sacrificed and BM cells was flashed from tibia/femur. The percentage of human CD45+ (hCD45) cells in engrafted murine bone marrow is shown after staining for hCD45 and analyzed by flow cytometry. Mann−Whitney U test with FDR correction for multiple hypothesis testing, *P<0.05; **P<0.005.

**Supplementary Fig. 4: VAF of *SRSF2* Mutations of CD45+ grafts from NSG mice injected with *SRSF2* Mut CD3- AML cells.** NSG mice were treated with RKI-1447 (50mg/kg) or vehicle for 3 weeks after 5-week transplantation of one to five millions of *SRSF2* Mut CD3- AML cells (i.f.). Mice were sacrificed and BM cells was flashed from tibia/femur. CD45+ cells from the grafts is isolated and the DNA of which is sequenced by targeted amplicon sequencing. VAF (variant allele frequency) is indicated with bar plot in each mouse.

**Supplementary Fig. 5: The percentage of CD19+, CD33+ and CD19-CD33- engraftment in the xenograft model.** The percentage of CD19+, CD33+ and CD19-CD33- human CD45+ (hCD45) cells in engrafted murine bone marrow with/without RKI-1447 treatment were measured with by flow cytometry to determine myeloid or multi lineage engraftment. **a**, healthy CD34+ sample. **b-d**, SRSF2 Mut AML samples, (**b**) #278788, (**c**)#830163, (**d**) #800667.

**Supplementary Fig. 6: The percentage of CD19+, CD33+ and CD19-CD33- engraftment in the xenograft model.** The percentage of CD19+, CD33+ and CD19-CD33- human CD45+ (hCD45) cells in engrafted murine bone marrow with/without RKI-1447 treatment were measured with by flow cytometry to determine myeloid or multi lineage engraftment. **a-c,** *SRSF2* mut AML samples. (**a**)#150532, (**b**) #209945, (**c**)#160141

**Supplementary Fig. 7: Hematotoxicity and Metabolic stability of RKI-1447. a**, 1000-2000 healthy CD34+ cells are cultured with Myeloid, Erythroid and Megakaryocytic lineage differentiation cytokines for 7 days, followed by addition of RKI-1447 (0.01, 0.04, 0.33, 1 and 3μM) for 48 hours (**see method**). Cell viability is detected with CellTiter-Glo assay. **b**, RKI-1447 (1 µM) was incubated with human liver microsomes (HLM) and NADPH at 37 degrees for a time course using thin-walled PCR tubes. Compounds are quantified by LC/MS. Data is normalized to percent of input compound in buffer without microsomes and time-course is plotted with Graphpad Prism. RKI-1447 metabolized in HLM with a half-life of 11.42 minutes. Positive control Diclofenac metabolized in HLM with a half-life of 16.45 minutes**.**

**Supplementary Fig. 8: hERG assay of RKI-1447 and differentiation of MOLM14 cells under RKI-1447. a**, the cardiotoxic effects measured by the blockage of the human ether-a-go-go-related gene(hERG) potassium channel by Predictor™ hERG Fluorescence Polarization Assay. The assay is performed with different concentration (0.01μM, 0.04μM, 0.33μM, 1μM and 3μM) of RKI-1447 and Cisapride (positive control). Y axis is polarization (mP) normalized (where full binding is 0, and no binding is 100%).**b-e**, Differentiation of *SRSF2* WT and Mut MOLM14 cells are detected with different surface markers (CD14, CD11b, CD15, CD66) by flow cytometry after 24hr addition of RKI-1447 (0.5μM). **f**, gene expression of neuthrophil differentiation markers (*AZU1*, *CTSG*, *ELANE*, *MPO*) of *SRSF2* Mut MOLM14 cells before and after 8 hours’ exposure to RKI-1447 (0. 5μM); **g**, gene expression of neuthrophil differentiation markers (*AZU1*, *CTSG*, *ELANE*, *MPO*) of *SRSF2* WT MOLM14 cells before and after 8 hours’ exposure to RKI-1447 (0. 5μM).

**Supplementary Fig. 9: Principal component analysis of protein expression before and after exposure to RKI-1447 of AML2 and MARIMO cells. a** Principal component analysis plot of protein expression of WT and *SRSF2* Mut AML2 cells before (0hr) and after 2 (2h) and 8 (8h) hours of exposure to RKI-1447 (0.5µM). **b**, Principal component analysis plot of protein expression of WT and *SRSF2* Mut MARIMO cells before (0hr) and after 2 (2h) and 8 (8h) hours of exposure to RKI-1447 (0.5µM).

**Supplementary Fig. 10: GSEA analysis of protein expression before and after exposure to RKI-1447.**

**a**, GSEA analysis of protein expression pre-ranked based on the LOG2 fold change between time before exposure and eight hours after exposure to RKI-1447 (0.5µM) on MOLM14 WT cells. Significant enriched pathways from the GSEA analysis included the Cell Cycle (NES=1.4365,
Nominal p-value=0.0223) and G2M checkpoint pathways (NES=1.427, Nominal p-value= 0.01007). (Left) KEGG, Kyoto Encyclopedia of Genes and Genomes. (Right) Hallmark glycolysis gene set. NES, normalized enrichment score. **b**, GSEA analysis of protein expression pre-ranked based on the LOG2 fold change between time before exposure and eight hours after exposure to RKI-1447 (0.5µM) on MARIMO *SRSF2* Mut cells. Significant enriched pathways from the GSEA analysis included the Cell Cycle (NES=1.376, Nominal p-value=0.046) and G2M checkpoint pathways (NES=1.425, Nominal p-value=0.016). (Left) KEGG, Kyoto Encyclopedia of Genes and Genomes. (Right) Hallmark glycolysis gene set. NES, normalized enrichment score. **c**, GSEA analysis of protein expression pre-ranked based on the LOG2 fold change between time before exposure and eight hours after exposure to RKI-1447 (0.5µM) on AML2 *SRSF2* Mut cells. Cell Cycle (NES=1.0797888, Nominal p-value =0.325) and G2M checkpoint pathways (NES=0.83830416, Nominal p-value=0.86292) were not significantly enriched. (Left) KEGG, Kyoto Encyclopedia of Genes and Genomes. (Right) Hallmark glycolysis gene set. NES, normalized enrichment score.

**Supplementary Fig. 11: Cell cycle analysis of MOLM14 WT cells before and after exposure to RKI-1447.** Cell cycle of MOLM14 WT before and after 2hr, 4hr, 6hr, and 8hr exposure to RKI-1447(0.5µM). Different cell cycles are indicated with different colors in the plot. T-test, *P<0.05; **P<0.005; ***P<0.0005.

**Supplementary Fig. 12: 3D rendering of nucleus of *SRSF2* WT and Mut MOLM14 cell lines before and after exposure to RKI-1447.** **a**, *SRSF2* WT and Mut MOLM14 cells, as indicated, were either left untreated or treated with RKI-1447 (0.5 µM) for 24 hours before fixation. Cells were labeled with DAPI to visualize nuclear morphology imaged on the Olympus confocal microscope as described in Materials and Methods, 3D volumes were subjected to 3D rendering using IMARIS software. Rendered surfaces are shown in dark blue. **b**, Morphometric characteristics of rendered volumes were extracted nuclear area was plotted in the box and whiskers format. Midline corresponds to median value, box includes central 50% of distribution, and whiskers correspond to upper and bottom 25%. Note, that bearing the mutation as well as treatment with RKI-1447 causes statistically significant increase in nuclear area. N= 114 for WT, N=126 for WT+RKI-1447, N=116 for Mut, N=119 for Mut+RKI-1447. The two-sample T-test with two-tailed distribution was used for evaluation of statistical significance. See also **Fig. 4a**.

**Supplementary Fig. 13: TEM of *SRSF2* WT and Mut MOLM14 cells before and after exposure to RKI-1447.** TEM, showing higher magnification images corresponding to the images shown in **Fig. 4b**. (**a** and **b**), lower magnification images with rectangles marking the enlarged areas, shown in **a**’ and **b**’, respectively. N and C mark nuclear and cytoplasmic areas, respectively. Arrows or arrowheads marked “Mt” point to microtubules, shown in longitudinal or cross sections, respectively. The arrow in a’-right points to the inter-lobular sheet, connecting the two nuclear segments. The twin-arrow in b’-left (MM) point to the attached nuclear membranes, and the arrow marked “L” point to the nuclear laminae.

**Supplementary Fig. 14: Confocal images of LaminB1** **and microtubules staining of *SRSF2* WT and Mut MOLM14 cells before and after exposure to RKI-1447.** *SRSF2* WT or Mut MOLM14 cells were either untreated or 24-hour treated with RKI-1447 (0.5 µM), as indicated. LaminB1 labeling shown in green and microtubules in red. **a**, Top to bottom rows: single confocal slices bottom to top direction extracted from 3D volume. The depth of the WT volume ranges from 0 to 11.4 μm, with confocal slices shown at intervals: 3.8 μm and 7.6 μm, the mut volume ranges from 0 to 12.6 μm with confocal image intervals at 4.2 μm and 8.4 μm, WT + RKI-1447 is 12 μm deep with image intervals 4 μm and 8 μm, and Mut + RKI-1447 has a z-depth of 11.1 μm with image intervals at 3.7 μm and 7.4 μm. These are presented in order to give the view of depth of nuclear indentations and degree of lobulation. See also Supplementary movie 1. **b**, selected enlarged insets from images presented in panel **a**. Cells in **a.** are outlined with dotted white line, and presented in panel **b**. according to numbers and treatment conditions. Note the presence of the microtubule organizing center at the base of nuclear deformation – indicated by bright microtubule labeling in first and third columns. Different focal planes reveal deep narrow indentations positive for LaminB1 and containing microtubules as indicated by red labeling (arrowheads in second and fourth columns).

**Supplementary Fig. 15: Z-stack confocal images of microtubules staining of *SRSF2* WT and Mut MOLM14 cells before and after exposure to RKI-1447**

*SRSF2* WT or Mut MOLM14 cells, as indicated, were either left untreated or treated RKI-1447 (0.5 µM) for 24 hours before fixation. Cells were labeled with anti-α-tubulin antibodies to label microtubules (shown in green). The z-stack images were and images were acquired using the Leica SP8 scanning confocal microscope (shown in green). The confocal slices are represented from top to bottom on the figure and range from top to bottom by the sample. The z volume was divided in six equal distances from the first slice chosen to the last. The slices are approximately 1.5 μm apart. Note, the prominence of microtubule bundles, especially in mutant cells upon RKI-1447 treatment. See also **Fig. 4f**.

**Supplementary Fig. 16: Dose response of Blebbistatin against *SRSF2* WT and Mut MOLM14 cell lines.** Viability of *SRSF2* WT and Mut MOLM14 cell lines were measured after 48 hours’ exposure to Blebbistatin at concentration of 0.037μM, 0.129μM, 0.369μM, 0.99 μM and 29.90μM.

**Supplementary Fig. 17: F-actin staining of *SRSF2* WT and Mut MOLM14 cells before and after exposure to RKI-1447.** *SRSF2* WT and Mut MOLM14 cells, as indicated, were either left untreated or treated with RKI-1447 (0.5µM), for 24 hours before fixation. **a**, Cells were labeled with fluorescent phalloidin to visualize actin (shown in red). 3D volumes were taken on the Leica SP8 scanning confocal microscope, represented slices corresponding to mid-plane of cells are shown. Clear exclusion of actin from nuclear region was observed in WT, but not in mutant cells. **b**, Quantification of the nuclear vs cytoplasmic actin was performed, with N values as follows N = 36 for WT, N= 26 for mut, N=29 for WT + RKI-1447 and N=26 for Mut + RKI-1447. Note, that the nuclear-to- cytoplasmic actin ratio was higher in mutant compared to WT, and higher in RKI-1447-treated WT cells, compared to non-treated WT nuclei.

**Supplementary Movie 1**

*SRSF2* WT and Mut MOLM14 cells, as indicated, were either left untreated or treated with 0.5 μm RKI-1447 for 24 hours before fixation. Cells were labeled with anti-LaminB1 antibodies to outline nuclear membrane (shown in green) and anti-α-tubulin antibodies to label microtubules (shown in red). 3D volumes were acquired on Olympus scanning confocal microscope. The slices were acquired as described in Materials and Methods with a z-step size of 0.3 μm and made into .avi movies starting at the bottom, and proceeding towards the top. Use the "Insert Citation" button to add citations to this document.
